# NeuroConText: Contrastive Learning for Neuroscience Meta-Analysis with Rich Text Representation

**DOI:** 10.1101/2025.05.23.655707

**Authors:** Fateme Ghayem, Raphaël Meudec, Jérôme Dockès, Bertrand Thirion, Demian Wassermann

## Abstract

Brain meta-analysis is the common way to gather information about human brain function across the existing literature in order to formulate hypotheses and contextualize new findings. However, automated meta-analysis tools face challenges such as inconsistent terminology and difficulties in analyzing long texts and capturing semantic meaning because they still rely on bag-of-words approaches; furthermore, sparse coordinate reporting in articles distorts the activation distribution due to incomplete data. This paper introduces NeuroConText, a predictive text-to-brain modeling framework designed to support brain meta-analysis by bridging neuroscience text, brain location coordinates, and brain images within a shared latent space. This framework follows the predictive brain meta-analysis paradigm: it learns a regression from text descriptions to whole-brain activation maps and also enables the retrieval of relevant studies through contrastive learning, optimizing a multi-objective loss that combines retrieval and reconstruction objectives. Furthermore, NeuroConText supports second-level statistical synthesis by providing activation associated with top-K retrieved studies that can serve as input to coordinate-based meta-analysis (CBMA) methods. NeuroConText also leverages large language models (LLMs) to capture neuroscientific information from full-text articles, plus an LLM-based text augmentation strategy to handle short-text inputs. Quantitative and qualitative analyses demonstrate NeuroConText’s ability to enhance text-to-brain retrieval performance and reconstruct brain maps from neuroscience texts. We also show that predictive brain meta-analysis tools can infer brain activations in regions discussed in articles but absent in reported coordinates, potentially addressing the challenge of sparse coordinate reporting.

## 1 Introduction

Hundreds of articles are published annually in the field of human neuroscience, highlighting the continuous accumulation of results in cognitive and clinical neuroscience. Manually collecting and analyzing information from such a large body of research is inefficient and thus requires automated methods (Yarkoni et al., 2011). Each study may examine different aspects of behavior, use specific methods, or emply unique experimental protocols. This adds layers of complexity to the integration of these diverse results into a coherent answer to a specific research question (Poldrack & Gorgolewski, 2014). Individual neuroscience studies are often challenged by small sample sizes and limited statistical power. This affects the reliability of the reported findings and reduces the likelihood that a statistically significant result reflects a true effect (Button et al., 2013). In addition, different framing within the literature leads to terminological inconsistencies; for example, different studies may use terms such as *cognitive decline* and *cognitive impairment* interchangeably (Poldrack & Yarkoni, 2016).

**Brain meta-analysis** is often used to produce a synthesis of the results reported in the literature to highlight the neural correlates of cognitive or neurological concepts (Yarkoni et al., 2011). It combines results from multiple studies, which improves the reliability of the associations between brain locations and cognitive or clinical concepts (Eickhoff et al., 2012). It serves three key purposes: first, it synthesizes information from the literature to summarize the state of knowledge in any subfield of human neuroscience. This allows researchers to see where the field stands collectively and highlight consensus or inconsistencies (P. T. Fox & Lancaster, 2002). Second, meta-analysis provides context to interpret new results by comparing novel experimental data with locations reported in the literature. This clarifies how new data align with or deviate from established findings (Wager & Smith, 2003). Third, meta-analysis generates hypotheses on candidate brain regions or relevant cognitive domains (Eickhoff et al., 2012).

In human neuroscience studies, experimental results are typically available in three modalities: the text of articles, brain activation coordinates reported in the tables of articles, and brain images available in public repositories at various levels of processing. Integrating these diverse data types is important, because they complement each other: texts give detail on cognitive hypotheses, cognitive conclusions, experimental procedures and provide cognitive concepts jointly with brain locations; brain coordinates tables provide a synthetic account of the main results of each publications, directly comparable across studies; brain images provide a more complete view of the correlates of cognition and diseases. Recent advances in large language models (LLMs) have demonstrated their ability to predict neuroscience research outcomes by synthesizing vast amounts of literature and making forward-looking predictions. BrainGPT is an LLM fine-tuned on neuroscientific publications to specialize in neuroscience research, and it has shown that LLMs can outperform human experts in predicting study results (Luo et al., 2024).

Bridging activation coordinates with brain images allows researchers to identify consistent activation patterns across studies (Eickhoff et al., 2012; M. D. Fox & Raichle, 2007; Laird, Fox, et al., 2005). Furthermore, connecting text from neuroscientific articles to brain activation coordinates can help to integrate textual data with imaging results for hypothesis generation and validating new results (Yarkoni & Westfall, 2017; Yarkoni et al., 2011). **Coordinate-based meta-analysis (CBMA)** is a meta-analysis tool that evaluates the stereotactic brain coordinates reported in neuroscience articles. It links neuroscience terms and descriptions to specific brain regions (Eickhoff et al., 2012). CBMA leverages the fact that virtually all recent neuroimaging studies report standardized stereotactic coordinates, typically in MNI or Talairach space; in practice, coordinates from different studies are converted into a common reference space, most often MNI, which enables aggregation across studies and gives absolute meaning to reported locations. (Collins et al., 1994; Mazziotta et al., 1995).

Several brain meta-analysis tools have been developed to link text and brain activation coordinates, ranging from manually curated databases to fully automated approaches (Laird, Lancaster, & Fox, 2005; Yarkoni et al., 2011). Traditional CBMA methods aggregate reported coordinates using voxel-wise statistical procedures, whereas more recent **predictive approaches** learn a direct mapping from text to whole-brain maps, bypassing coordinate-level statistical tests (Dockès et al., 2020; Ngo et al., 2021). Here we use the term “predictive” to distinguish between regression-based models that directly estimate brain maps from text and CBMA methods that focus on coordinate-level statistical inference. We provide a detailed overview of these methods and their limitations in Section 2.

Brain meta-analysis tools can help researchers navigate the rapidly expanding neuroimaging literature efficiently and allow them to extract relevant information for hypothesis testing and knowledge discovery. Retrieval and reconstruction are two key processes that support these goals, both involving the translation of textual descriptions into spatial brain data. This facilitates the synthesis of new findings and generates new insights into the neural activations that underlie cognition and behavior. The **retrieval** task focuses on associating relevant brain locations or images with specific textual queries or hypotheses, a process central to hypothesis testing and literature review. For example, when researchers search for studies related to a particular cognitive function, *e*.*g. working memory*, they retrieve the brain coordinates associated with that function. This enables the integration of findings across diverse studies and supports the development of new hypotheses. On the other hand, the **reconstruction task** involves mapping textual descriptions of brain activity (e.g., from published studies) onto spatial brain coordinates or activation patterns. This process is essential for interpreting and aggregating findings across studies, allowing the creation of meta-analytic maps that summarize brain-behavior relationships. Reconstruction generates brain maps representing the spatial distribution of activations associated with a given textual description, such as a cognitive function or experimental condition.

Existing brain meta-analysis tools are optimized for reconstruction and fall short when it comes to retrieval. A core challenge in the retrieval task lies in accurately aligning textual descriptions with spatial brain representations. **Contrastive learning** offers a promising solution for associating (i.e. retrieving) an input text with its corresponding image. This family of methods effectively bridges text and image by establishing a shared latent space between the two modalities (He et al., 2020; Radford et al., 2021; Tian et al., 2020). **CLIP** is a key example that exploits contrastive learning (Radford et al., 2021). Contrastive modeling learns to identify correct (image, text) pairings from a set of potential matches by optimizing cosine similarity for true pairs and minimizing it for mismatched pairs. This suggests that CLIP has the potential to effectively improve retrieval between texts and their corresponding MNI coordinates.

In this paper, we introduce **NeuroConText**, a meta-analysis model designed to address the retrieval task in brain meta-analysis (Meudec et al., 2024). NeuroConText captures the articles semantics with an LLM and associates them with the reported brain locations. A preliminary short publication of the same work in (Meudec et al., 2024) demonstrated the promising retrieval performance of NeuroConText. Here, we expand NeuroConText and introduce a *joint text-to-brain retrieval and reconstruction* framework with dual training objectives which leverage contrastive and mean squared error (MSE) losses to address retrieval and reconstruction tasks simultaneously. We also extend our experiments and provide extensive quantitative and qualitative analyses for retrieval and reconstruction tasks on left-out articles, and demonstrate NeuroConText’s ability to bridge text and brain activation. We further enhance NeuroConText’s performance and generalization to short-text inputs by introducing *LLM-based text augmentation*.

The remainder of the paper is organized in four parts: Section 2 reviews related work and places our study in context. Section 3 details the NeuroConText methodology, clarifies the workflow, and provides the specifications needed for reproducibility, while Section 4 presents the brain meta-analysis and neuroscientific results, which are further interpreted in the Discussion Section 5. More specifically, we introduce data preparation and preprocessing of the articles’ activation coordinates and text in Section 3.1. NeuroConText framework combining contrastive and MSE losses is introduced in Section 3.2. The inference task for reconstruction and the downstream task for retrieval are explained in Sections 3.3. We evaluate NeuroConText’s ability to retrieve brain maps from text through quantitative and qualitative assessments in Section 4.2. We demonstrate NeuroConText’s capability to reconstruct brain maps from text using quantitative and qualitative methods in Section 4.3. To enhance NeuroConText’s performance with short text inputs, we leverage LLMs for the augmentation of short-text inputs detailed in Section 3.4, with the results detailed in Section 4.4. Finally, Section 5 discusses the outcomes of this paper.

## 2 Prior Work

Early efforts for CBMA, such as **BrainMap** (Laird, Lancaster, & Fox, 2005), provide structured information about published neuroimaging studies by curating reported brain coordinates and annotating them according to a specialized taxonomy. However, as the neuroimaging literature grows rapidly, manual curation of databases is not scalable enough to cover all publications. To address scalability, **NeuroSynth** (Yarkoni et al., 2011) automates the meta-analysis of neuroscientific studies by identifying relevant articles through the terms in their *abstracts* and extracting brain activation coordinates to create brain statistical maps. In NeuroSynth, texts are represented by the frequency with which they appear in the abstract of an article. It uses multilevel kernel density analysis (MKDA) (Wager et al., 2009) to convert reported coordinates into statistical maps. This automated process allows efficient and automated analysis of the entire corpus. However, NeuroSynth has some key limitations. It relies on statistical testing, so it is not designed to accurately predict novel queries. It also uses a bag-of-words approach based only on abstracts, not full texts, so it may miss complex topics, new research terms, and syntactic information, such as negations and conditionals. Finally, it struggles with rare terms, which limits how well it can cover less common research topics. Building on this, **NeuroQuery** (Dockès et al., 2020) introduces a **predictive** brain meta-analysis model that directly estimates brain maps from text, addresses several of these limitations by using a multivariate model trained on the *full texts* of publications rather than just abstracts to predict relevant brain locations for neuroscience queries. It uses kernel density estimation (KDE) (Scott, 2015; Silverman, 2018) to represent lists of coordinates as brain maps. NeuroQuery employs the term frequency-inverse document frequency (TF-IDF) method (Salton & Buckley, 1988) to analyze texts, measuring the importance of words within a document relative to their frequency across the entire corpus. It applies semantic smoothing techniques that leverage term co-occurrences to improve the identification and mapping of infrequent terms. Although NeuroQuery improves upon NeuroSynth by using full texts and a broader vocabulary, it remains constrained by its reliance on a bag-of-words model that does not fully take into account the semantics of the text, treating word occurrences independently without considering their context or relationships. More recently, **Text2Brain** (Ngo et al., 2021) has introduced a large language model (LLM)-based *predictive* approach to link text with reported brain locations. It uses a transformer-based text encoder built on the SciBERT architecture (Beltagy et al., 2019) to represent the semantics of the text. During training, Text2Brain uses an augmentation strategy where it selects various types of text inputs, including the first sentence of table captions, articles’ titles, keywords, abstracts, and randomly selected subsets of sentences from the discussion sections. The encoded text is then fed into a 3D convolutional neural network (CNN) to estimate spatial brain activation maps.

Although previous tools have advanced automated meta-analysis by linking text to the brain, they are limited by weak text representations, inadequate retrieval capabilities, and restricted input formats. **NeuroConText** is a predictive model that addresses these challenges by using contrastive learning on full-text data to enable both retrieval and reconstruction of brain activation patterns simultaneously. In the next section, we detail its architecture and learning strategy.

## 3 Methods

In this section, we describe **NeuroConText**, a joint text-brain retrieval and reconstruction model based on contrastive learning and the full semantic context of neuroscience texts. NeuroConText links neuroscience texts and brain activation peak distribution maps by learning latent representations of text and brain locations. These representations support the retrieval of brain activation patterns from text and the estimation of relevant brain maps from neuroscience texts.

NeuroConText consists of four main modules: 1) data preparation and preprocessing, 2) training, 3) inference and downstream tasks, and 4) text augmentation at the inference phase. In the following, we detail each module.

### Ethics Statement

This study did not involve the collection of new data from human participants. All analyses were conducted on publicly available datasets. Therefore, no institutional ethics approval or informed consent was required.

### 3.1 Data Preparation and Preprocessing

In this section, we describe the procedures that we use for the preparation of text and coordinates data in the neuroscientific articles. We focus on transforming full-body texts and reported brain coordinates into representations suitable for NeuroConText’s computational analysis.

- **Text data representation using LLM embeddings:** To capture the semantics of neuroscientific articles, we use LLMs that leverage attention mechanisms. We aim to process the **entire** text of each article. However, due to the token size limitations of LLMs, we segment the full-body text into fixed-size chunks. Specifically, we use Mistral-7B (Jiang et al., 2023) with a context window of 8,192 tokens, which ensures that each chunk does not exceed this limit. For texts shorter than the chunk size, we apply padding using a special token to maintain uniform input dimensions. After obtaining the embeddings for each chunk, we aggregate them through mean pooling to create a unified text representation for each article. This approach ensures that the semantic information from the entire article is preserved while respecting the token constraints of the LLM.
- **Representing coordinates as brain maps using KDE:** We represent reported coordinates as smooth brain maps using kernel density estimation (KDE) where each peak is convolved with a Gaussian kernel and the contributions are summed to obtain a continuous 3D map. This construction is similar to the activation maps used in ALE-based CBMA methods, where reported foci are transformed into smooth spatial activation maps (Rubin et al., 2017; Scott, 2015; Silverman, 2018). So, we proceed as follows: Let **G** = [**g**_1_, …, **g**_*v*_] ∈ ℝ^3*×v*^ represent a fixed sampling grid of voxel positions in a 3D space with the total number of *v* voxels, where each **g**_*j*_ ∈ ℝ^3^ denotes the position of the *j*-th voxel. Let **P** = {**p**_1_,, **p**_*l*_} be the set of peak coordinates extracted from the brain image, where each **p**_*i*_ ∈ ℝ^3^ is the position of the *i*-th peak, and *l* is the number of peaks. The RBF kernel *κ*_*σ*_ centered at each peak is defined by:

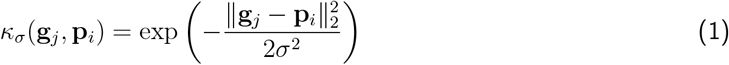

where *σ* is the kernel parameter. Then, the KDE representation of the estimated image **m** = [*m*_1_, …, *m*_*v*_]^*T*^ ∈ ℝ^*v*^ over the grid **G** is calculated by summing the contributions of the Gaussian kernels centered at each peak across all grid points:

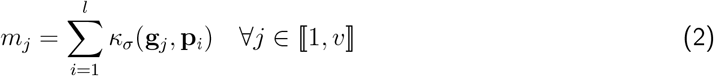 This equation defines how each voxel **g**_*j*_ in the grid is influenced by the Gaussian kernels placed at each peak **p**_*i*_, which results in a continuous map **m** proportional to the spatial density of activity peaks.
- **Brain image dimension reduction with DiFuMo atlas:** To address the high dimensionality of the brain maps reconstructed by KDE, we employ the Dictionary of Functional Modes (DiFuMo), a probabilistic atlas that effectively reduces data dimensionality (Dadi et al., 2020). This method provides a linear decomposition of the brain maps, where the data matrix **M** ∈ ℝ^*v×n*^, containing *n* brain maps, is represented as **M** = **DX**. Here, **D** ∈ ℝ^*v×k*^ is the dictionary matrix containing basis signals called atoms as its columns, under non-negativity and sparsity constraints. The coefficients matrix **X** ∈ ℝ^*k×n*^ contains the linear combination coefficients for this decomposition. The number of atoms *k* = 512 is significantly smaller than the number of voxels *v*, which ensures a substantial dimensionality reduction. In this study, we use the set of *k* DiFuMo coefficients as image features. The process of deriving DiFuMo coefficients from KDE maps is formulated as a linear regression problem.

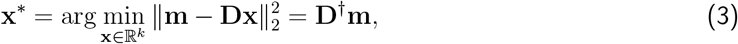

where **D**^*†*^ = (**D**^*T*^ **D**)^−1^**D**^*T*^ is the pseudo-inverse of **D**.

### 3.2 Training module

To jointly model the relationship between neuroscience text and brain activation patterns, NeuroConText model is trained using a contrastive learning objective that aligns representations of text and brain maps in a shared latent space. The model learns to associate semantically related pairs while distinguishing them from unrelated ones. This training process enables both retrieval (matching text to brain data) and reconstruction (generating brain maps from text) within a unified framework. Fig.1 shows NeuroConText’s training process, which consists of the following steps:

- **Encoding and Decoding:** To perform the retrieval task, we define a shared latent space by mapping text embeddings through a text encoder and DiFuMo representations of brain maps through an image encoder. We use **Mistral-7B** (Jiang et al., 2023) as the LLM to generate text feature representations with an embedding dimension *d* = 4096. The latent space dimension is set to *k*, and a bottleneck in the text encoder reduces the dimension from *d* to *k*. Next, we implement a decoder to reconstruct DiFuMo coefficients of brain maps from the text’s latent representation. The architecture of the text encoder, image encoder, and decoder are depicted at the bottom of Fig.1. The encoders and decoder consist of blocks, each containing a fully connected (FC) layer with *k* units, followed by GELU activation, dropout (50%), and layer normalization. Specifically, the text encoder comprises three blocks, with the first block including a residual connection. The image encoder contains three blocks, and the decoder has two blocks.
- **Multi-objective loss and alternating optimization:** We train **NeuroConText** to perform joint text-brain retrieval and brain map reconstruction. Consider Θ = [*θ*_text_, *θ*_img_, *θ*_dec_] to be NeuroConText’s trainable model parameters corresponding to the weights of the text encoder denoted by the function *f*_text_ : ℝ^*d*^ → ℝ^*k*^, image encoder *f*_img_ : ℝ^*k*^ → ℝ^*k*^, and decoder *f*_dec_ : ℝ^*k*^ → ℝ^*k*^ with the architecture shown at the bottom of Fig.1 detailed in the previous part. *k* = 512 is the DiFuMo dimension, and *d* = 4096 is the Mistral-7B embedding size. We show the input text embeddings of *N* samples by the matrix **Y** = [**y**^[1]^, …, **y**^[*N*]^] where columns **y**^[*i*]^ ∈ ℝ^*d*^ represent the *i*^th^ sample. We denote the DiFuMo coefficients of the input data by the matrix **X** = [**x**^[1]^, …, **x**^[*N*]^] where each column **x**^[*i*]^ ∈ ℝ^*k*^ corresponds to the sample *i*. NeuroConText’s model optimizes a multi-objective loss function that minimizes the sum of the InfoNCE contrastive loss for retrieval and the mean squared error (MSE) loss for reconstruction:

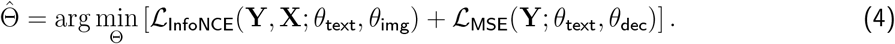 Here, ℒ_InfoNCE_ is the InfoNCE loss, defined as:

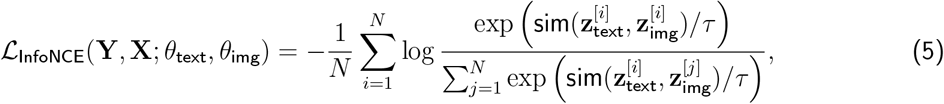

where 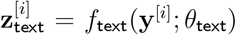 and 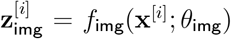 are the latent representations of the text and image, respectively. The function 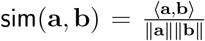 denotes the cosine similarity, with ⟨·, ·⟩ as the inner product and ∥ · ∥ as the Euclidean norm, and *τ* is a temperature parameter. This loss encourages alignment of the text and image representations in the shared latent space. The second objective function, ℒ_MSE_, is the MSE loss, defined as:

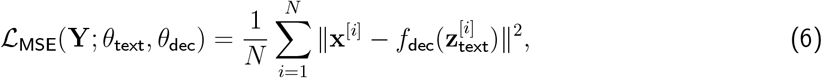

**Figure 1.**
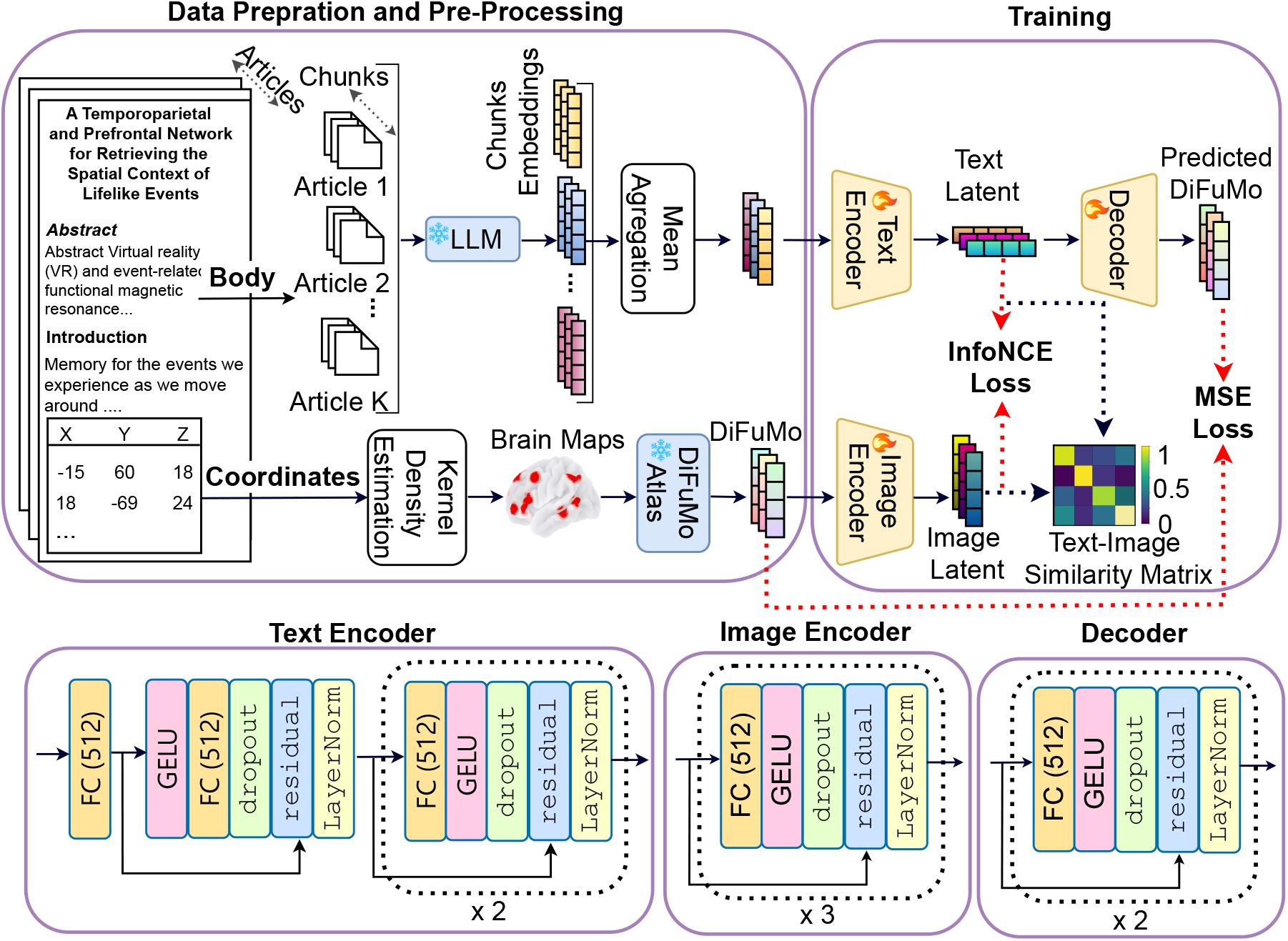
NeuroConText training : The training pipeline consists of three main steps: (1) Data preparation and preprocessing: Neuroscience articles are split into chunks, and a pretrained large language model (LLM) is used to encode each chunk and generate LLM-based embeddings. These embeddings are aggregated using mean aggregation. In parallel, brain coordinates from the articles are transformed into brain maps using kernel density estimation (KDE), which are then encoded in the DiFuMo atlas. (2) Encoding into a shared latent space and decoding: Text embeddings are passed through a text encoder, while DiFuMo coefficients are passed through an image encoder, both of which are trained to project inputs into a shared latent space. The architecture of the text encoder, image encoder, and decoder is shown below. (3) Learning objectives: A contrastive loss using the InfoNCE objective ensures that text and brain map embeddings from the same article are brought close in the latent space. A mean squared error (MSE) loss guides the decoder to reconstruct brain maps in the DiFuMo space from the text embeddings. In this figure, the blue modules with the snowflake icons and the yellow modules with the fire icons denote the frozen (pretrained) and trainable models, respectively.

Learnable parameters Θ = [*θ*_text_, *θ*_img_, *θ*_dec_] for the text encoder, image encoder, and the decoder respectively, are structured as matrices of dimensions *k × k* for each FC layer. This configuration involves approximately 4.2 million parameters.

For training NeuroConText’s model parameters Θ, we first optimize the parameters *θ*_text_ and *θ*_img_ of the encoders to minimize the InfoNCE loss to ensure alignment between text and image representations in the shared latent space. We optimize *θ*_text_ and *θ*_dec_ to minimize the MSE loss by focusing on the reconstruction of DiFuMo coefficients from text embeddings. We alternate between these two optimization steps, wherein during each iteration, *θ*_text_ is adjusted based on the contrastive loss initially and later fine-tuned using the MSE loss.^1^

### 3.3 Inference Module and Downstream Task

Once NeuroConText’s model has been trained, we leverage the model to perform inference and consider a downstream task to evaluate the efficiency of our model. This is illustrated in Fig.2.

- **Text-brain retrieval (downstream task):** We perform a retrieval task to identify the relevant brain map for a given textual query. As depicted in Fig. 2-bottom, we compare the input text embedding with brain map embeddings from our dataset in the shared latent space. We calculate a text-image similarity vector and rank the brain map embeddings to determine if the ground-truth map is among the top *K* retrieved items.
- **Brain map reconstruction (inference):** The inference task focuses on reconstructing a brain map from a given text. NeuroConText provides two complementary reconstruction modes: (i) a predictive map decoded directly from text, and (ii) a statistical map similar to CBMA which is computed directly from the KDE maps. For the predictive map, as shown in Fig.2-top, we input text into the LLM to generate chunk-level embeddings. These embeddings are aggregated to form the input text representation. This representation is passed through the trained text encoder and decoder to produce a brain map in the DiFuMo space. We then apply the DiFuMo inverse transform to recover the brain map in the original image domain. In addition, for the CBMA-based reconstruction method, we perform a voxel-wise second-level statistical analysis comparing the KDE maps of the top-*K* retrieved studies against those of the non-retrieved studies. This results in a NeuroConText statistical map that highlights significant differences in activation patterns between the two groups.

**Figure 2.**
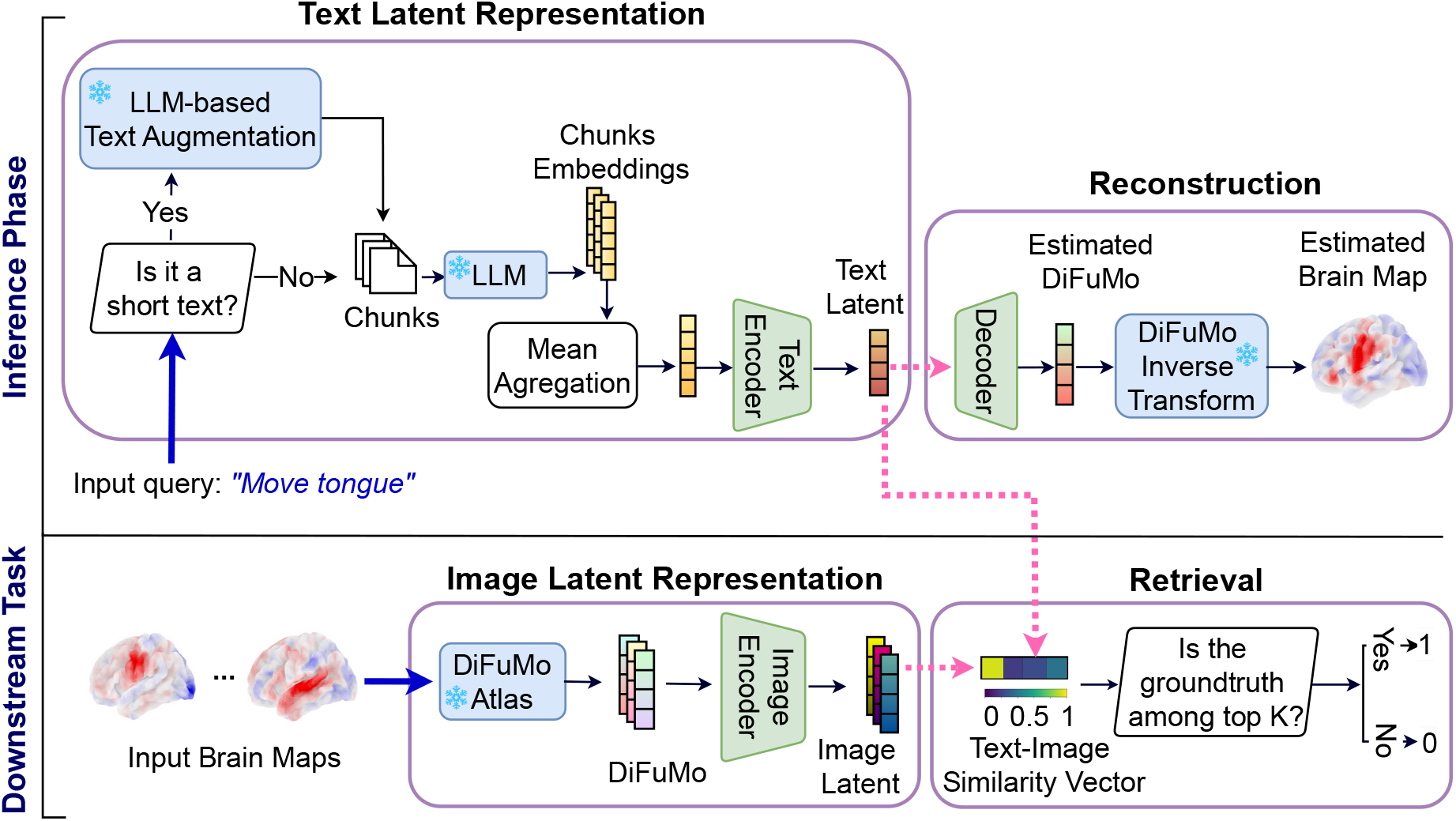
Use of NeuroConText-Inference phase (Top): The inference pipeline includes two tasks: brain reconstruction from an input text and article retrieval. Short input queries (e.g., “Move tongue”) are processed through LLM-based text augmentation, then the augmented text is split into chunks. Next, the chunks are encoded and aggregated to form the input text embedding. This aggregated embedding is passed through NeuroConText’s trained text encoder and decoder (green modules) to reconstruct the DiFuMo coefficients corresponding to the KDE representation of the article’s reported coordinates. Then, we map estimated DiFuMo coefficients into the voxel space using the DiFuMo inverse transform. **Downstream Task (Bottom):** In the downstream task, we retrieve NeuroVault’s brain contrast maps corresponding to the input queries, which are NeuroVault’s short-text contrast definitions. To do so, the aggregated input text embedding is compared to the brain map embeddings in the database in the latent space through the text-image cosine similarity vector, where we expect to retrieve the corresponding map among the top-k retrieved items. In this figure, the blue modules with the snowflake icons and the green modules denote the frozen (pretrained) and trained NeuroConText models, respectively.

### 3.4 LLM-based text augmentation for short input text at inference

While NeuroConText is trained on full-text articles, we adapt it for use with small texts, such as functional task descriptions or contrast specifications, similar to those available in the Cognitive Atlas. We note that the model is trained only on full-text articles, and this adaptation is done at inference time by expanding short inputs via LLM-based text augmentation, without any additional fine-tuning on a short-text corpus. This can result in suboptimal performance due to a distribution shift in the embeddings generated by the language model for short and long texts, that is, the statistical properties of the embeddings differ depending on text length^2^.

To enhance NeuroConText’s ability to reconstruct brain maps from short-text descriptions, we employ LLM-based text augmentation using GPT-4o. The augmentation process expands short descriptions into long article-like formats by including sections such as introduction, methodology, experiments, and discussion. To ensure the validity of the augmented texts and minimize hallucination risks, we ask domain experts to review the results, though occasional discrepancies may still exist. In the Results Section, we evaluate the impact of this distribution shift by comparing NeuroConText’s performance on NeuroVault (Gorgolewski et al., 2015) short-text inputs before and after text augmentation.^3^

### 3.5 Evaluation metrics

We consider different evaluation metrics for text-brain retrieval and reconstruction tasks.

- **Retrieval evaluation metrics:** To evaluate how well the model can retrieve the correct brain map given a textual description, we use similarity scores (as illustrated in Fig. 2) between the input text and all candidate brain maps in the test set. The evaluation assesses whether the model ranks the correct brain map higher than the incorrect ones. In this regard, we use the following two metrics:

#### Recall@K

This metric measures how often the correct brain map is found among the top-K most similar maps retrieved by the model for each input text. A higher Recall@K indicates that the model better aligns text inputs with their corresponding brain maps.

#### Mix&Match

This metric, which is adapted from Mitchell et al. (Mitchell et al., 2008), tests whether the model can correctly identify the brain map that matches the input text. For each text, it compares the similarity of the retrieved map to the ground truth map versus a randomly selected incorrect map. The score reflects how often the retrieved map is more similar to the ground truth map than the random map, with higher values indicating better retrieval accuracy.

- **Reconstruction evaluation metric:** We use **Dice Score**, as in the Text2Brain paper, to asses NeuroConText’s performance to predict the density of activation peaks from input text descriptions. The Dice Score measures spatial overlap between the predicted maps and actual activation maps. For a given threshold *t*, the Dice score *D*_*t*_ quantifies how well activations co-occur between the ground truth activation map **y** and the predicted map **ŷ**:

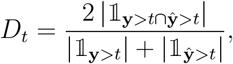

where 𝟙_**y***>t*_ is an indicator function that identifies voxels with activation values above the threshold *t*, and | · | represents the volume of activated regions in the image space.

Combining the above metrics, our evaluations assess NeuroConText’s ability to retrieve and generate brain maps from textual descriptions. This dual evaluation strategy reflects the model’s ability to bridge neuroscience text and brain maps.

### 3.6 Implementation Details and Code Availability

For the KDE representation of the coordinates, we set FWHM = 9*mm*. NeuroConText model was trained using a batch size of 125, with the Adam optimizer (Kingma & Ba, 2014). The learning rate for the encoder and decoder was set to 1 *×* 10^−4^, with a weight decay of 0.1. Dropout was applied at a rate of 0.6 for both the encoder and decoder to mitigate overfitting. The temperature parameter *τ* for the InfoNCE loss is set to 10. The training was performed over 50 epochs.

We used Python 3.10.9, PyTorch 2.0.1+cu117, and the Hugging Face Transformers library (version 4.39.3) for model implementation and inference. For text representation, we used the Mistral-7B-v0.1 model from the Hugging Face Model Hub, along with its associated tokenizer. To handle short input texts during inference, we employed LLM-based text augmentation using GPT-4o. For extracting Mistral embeddings of the text, we used a computing system consisting of a CPU node with 2 × Intel(R) Xeon(R) CPU E5-2660 v2 @ 2.20GHz, 256 GB RAM, and a GPU node with 3 × Nvidia V100 32G. For model training, inference, and other experiments, we conducted our work on a system equipped with a Tesla V100-DGXS GPU (Driver Version: 450.51.05, CUDA Version: 11.0) and an Intel(R) Xeon(R) CPU E5-2698 v4 @ 2.20GHz with 40 cores and 251 GB of RAM. The code for reproducing our results is publicly available at https://github.com/ghayem/NeuroConText. This repository includes preprocessed text and coordinates data and scripts for NeuroConText model training and inference.

## 4 Experiments and Results

### 4.1 Data Preparation and Experiment Setup

We downloaded and prepared neuroscience articles from PubMed Central (PMC) using Pubget. Pubget is an open access tool that extracts text, metadata, stereotactic coordinates, and Term Frequency-Inverse Document Frequency (TF-IDF) data from the articles (Salton & Buckley, 1988). We extended the NeuroQuery dataset (Dockès et al., 2020) by increasing its size from 14K to 20K articles, including more recent publications. After downloading the articles and obtaining their text and coordinates, we obtained the DiFuMo representation of the coordinates and the LLM embeddings of the full-body text. We split the neuroscience articles into 19K training and 1K test samples. We then trained the NeuroConText model with the training set and evaluated its performance in the retrieval and reconstruction tasks on the test set. We used a 19-fold cross-validation for model selection embedded in a 15-fold cross-validation to evaluate the model’s performance on unseen test data. We compared NeuroConText to two baselines: NeuroQuery and Text2Brain. For a fair comparison with the baselines and to avoid leakage, we ensure that articles used in NeuroQuery’s training set are excluded from the test set.

### 4.2 Brain Map Retrieval from Textual Input

In the first experiment, we evaluated the ability of NeuroConText to associate the text of neuroscientific articles with their corresponding coordinates reported in the articles. To assess NeuroConText’s performance on the retrieval task, we followed the procedure outlined in Section 3.3 and shown in Fig.2. We aligned the input articles’ text (title, abstract, full-body) with their corresponding coordinates set based on texts-coordinates cosine similarities in NeuroConText’s latent space. We then computed the retrieval metrics—Recall@K for *K* ∈ {10, 100} and Mix&Match—on the left-out articles according to their definitions provided in Section 3.5. To compare with other meta-analysis tools, we also calculated these metrics for NeuroQuery and Text2Brain using their pre-trained models to estimate brain maps from the article’s text in the test set and calculate their similarities with the KDE representation of the coordinates.

#### 4.2.1 NeuroConText leads in retrieving brain activation from long text inputs

Our experiments on brain map retrieval from text show that NeuroConText outperforms NeuroQuery and Text2Brain. When evaluating recall@10 for full-body text, NeuroConText achieves a score of 22.6%, far from NeuroQuery (7%) and Text2Brain (1.4%), as shown in Fig.3. This trend holds when the input text is the article’s abstract or title. Furthermore, in NeuroConText, the results for full-body text are higher compared to the abstract, with Recall@10 improving from 12.8% (abstract) to 22.6% (body text). This shows the impact and importance of leveraging the richer information in the full-body text compared to the relatively concise abstract.

**Figure 3.**
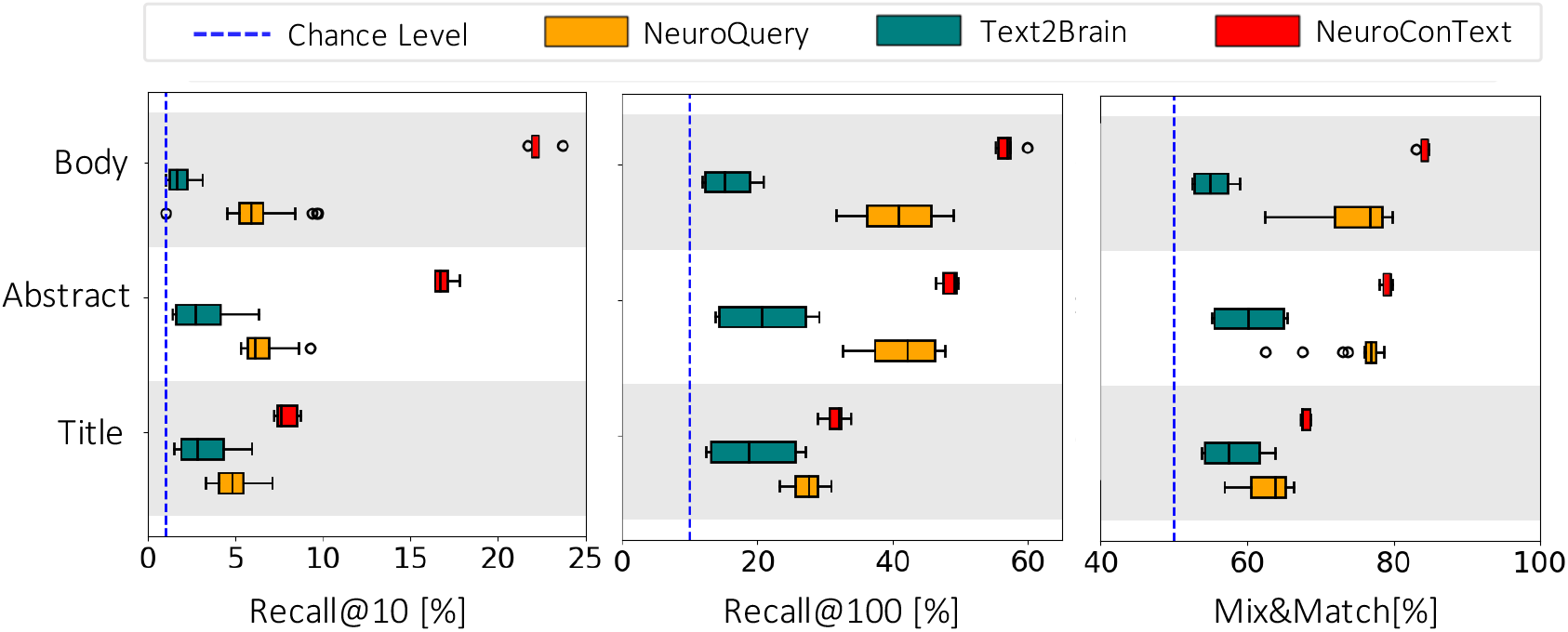
Retrieval of brain coordinates from articles’ text input: NeuroConText achieves higher retrieval scores compared to NeuroQuery and Text2Brain, particularly for full-body text input. This figure presents results on left-out articles: Given a text query—either the title, abstract, or body of a neuroscientific publication—we report Recall@10, Recall@100, and Mix&Match, computed based on cosine similarities between text and coordinate representations. The dashed blue line is the chance level used as a baseline. These metrics assess the ability to retrieve the coordinates reported in the articles that best represent the input text.

### 4.3 Reconstructing Brain Maps from Article’s Text

We assessed NeuroConText’s ability to generate brain activation maps from text latent representations to obtain insights into its internal latent representations. In this experiment, we used the full text of the articles as input. We compared NeuroConText’s reconstructed maps with the KDE representation of coordinates using the Dice Score metric explained in Section 3.5. For comparison with the baselines, we applied NeuroQuery and Text2Brain to the same test articles and computed Dice Scores between their reconstructed maps and the articles’ KDE representations.

#### 4.3.1 NeuroConText is on par with the state of the art in the reconstruction task for left-out articles

NeuroConText consistently achieves Dice scores on par with NeuroQuery and Text2Brain across all thresholds for brain map reconstruction from text, as shown quantitatively in Fig. 4. We also provide a qualitative comparison of the brain maps reconstructed by NeuroConText and the baseline models in Fig.5 for the best, median, and worst reconstruction performances of NeuroConText. Using the Mistral language model, we provide a summary of the articles and the brain regions discussed in the articles to compare the reconstructed maps with the content of the articles’ text. Notably, as shown in Fig. 5-a, these case studies include not only task-based cognitive neuroimaging articles but also structural and clinical neuroscience studies, illustrating that NeuroConText can reconstruct brain maps across diverse study types. Comparing the NeuroConText reconstruction with the KDE map in Article 1, we observe activation in the superior temporal gyrus in both maps, which also appears among the regions extracted from the article text. In Addition, activations are present in the angular gyrus, supramarginal gyrus, and insula—regions listed in the article, athough they are not observed in the KDE representation. In Article 2, we observe activations in the precentral gyrus, corpus callosum, and external capsules for NeuroConText, while Text2Brain shows additional activations, such as in the supplementary motor cortex. In the worst case for NeuroConText, Article 3, we observe overlaps with the KDE in the left insula and left supramarginal gyrus. In addition, for both NeuroConText and Text2Brain, activations appear in the left precentral gyrus and cerebellum VI, which are not explicitly present in the KDE map but are listed among the brain regions of the article.

**Figure 4.**
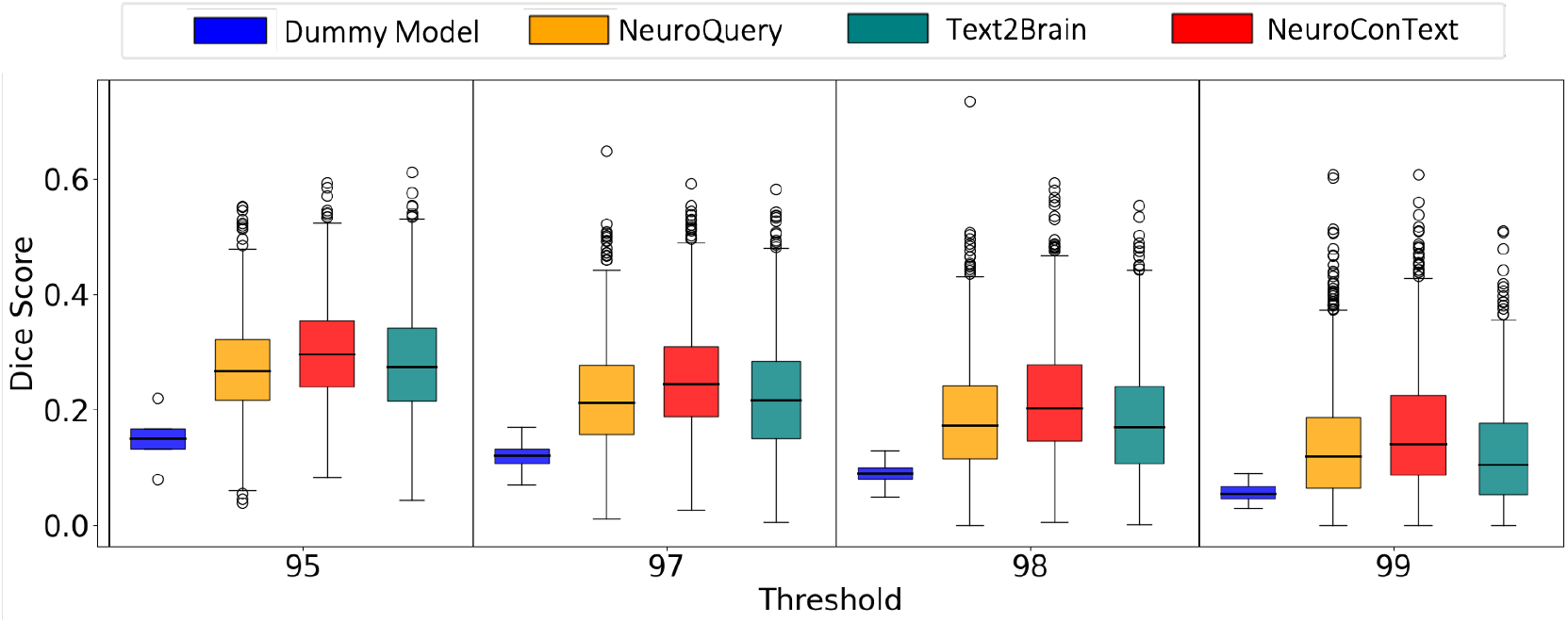
Reconstructing brain maps from articles’ text input: NeuroConText performs on par with NeuroQuery and Text2Brain in reconstructing brain activation maps from textual input. NeuroConText achieves comparable Dice scores across different thresholds. The thresholds correspond to percentiles of the brain maps histograms. This figure presents the Dice score evaluation of brain maps reconstructed from full-body text input using NeuroConText, NeuroQuery, and Text2Brain. The Dice Score measures the spatial overlap between the reconstructed maps and the KDE representation of the articles’ coordinates. The Dummy Model in blue serves as a baseline to highlight the added value of CBMA tools. Threshold values correspond to quantiles of the image histogram.

**Figure 5.**
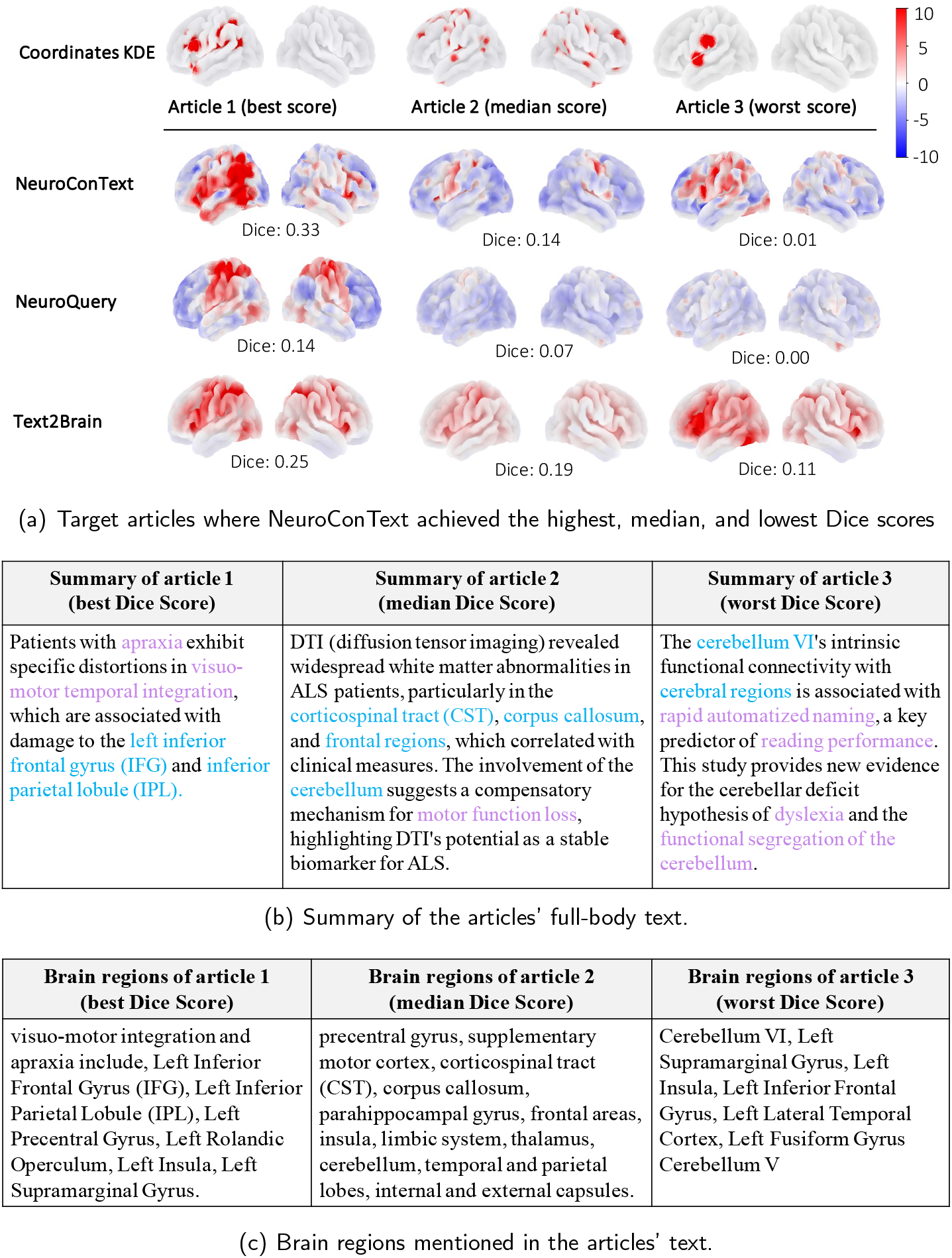
Qualitative comparison of reconstructed brain maps from articles’ text input: In the best-performing cases, NeuroConText performs on par with Text2Brain and NeuroQuery. In the median and worst cases, although the overlap with the KDE representation is smaller, we still observe that NeuroConText and Text2Brain reconstructions match with the brain regions listed for the articles. This suggests that these tools can infer activations beyond the sparse coordinates reported in the articles and highlights the limitations of using Dice scores based on KDE maps as the sole evaluation metric. (a) Comparison of brain activation maps reconstructed by different models for the articles with the best, median and worst achieved Dice scores. The colorbar represents the normalized intensity values of brain activation. (b) Summary of the articles. (c) Brain regions discussed in the articles.

While the main focus of this paper is on predictive brain map reconstruction from text, we also explored a complementary CBMA-based reconstruction approach. Specifically, we performed voxel-wise second-level statistical analysis between the top-200 retrieved studies and the non-retrieved ones, based on their KDE maps. Notably, the statistical maps show considerable spatial overlap with the predictive maps, yet the predictive maps tend to be more focal. A qualitative comparison between the predictive and statistical reconstructions is presented in Appendix 6.1.

### 4.4 LLM-Based Text Augmentation for Short Texts

To assess NeuroConText’s generalization to short-text inputs, we evaluate it on 196 contrast descriptions from the collection 2138 of the NeuroVault dataset (Pinho et al., 2018), along with some associated contrast maps. The ID of the chosen samples is provided in NeuroConText GitHub repository. NeuroVault is a large open-access repository of fMRI statistical maps from group or single-subject studies. Each NeuroVault sample includes associated cognitive terms or labels. It therefore provides a validation of NeuroConText’s ability to generalize from coordinate-based training data to real-valued statistical maps. A key challenge in this context is the **distribution shift** between LLM embeddings of long and short texts. By applying Principal Component Analysis (PCA) to text embeddings, we see a prominent separation between short NeuroVault contrast definitions and longer publication texts (abstracts and full-body) in the embedding space, along two principal components^4^. This suggests that the disparity in text length impacts how text information is represented and influences NeuroConText’s performance.

In this subsection, to quantitatively assess the impact of text augmentation, we apply NeuroConText to reconstruct brain maps from NeuroVault descriptions in two forms: (1) the original short descriptions and (2) augmented descriptions transformed into longer, article-like formats using GPT-4o. We evaluate the reconstruction accuracy using Dice scores over thresholds corresponding to the 95%, 97%, 98%, and 99% quantiles of the image to compare the performance of NeuroConText, NeuroQuery, and Text2Brain and to show the ability of each model to recover activation patterns from textual input. Next, we perform a qualitative analysis of brain map reconstructions to further evaluate the impact of text augmentation. We also study the impact of LLM-based text augmentation on retrieval performance. In the following, we describe the results.

#### 4.4.1 LLM-based text augmentation enhances NeuroConText reconstruction performance on short-text input

Table 1 shows the impact of text augmentation on reconstruction performance by comparing the results before and after the augmentation explained in Section 3.4 for the NeuroVault short-text descriptions. We evaluated the performance across various brain map thresholds. We see that augmented text significantly improves NeuroConText’s performance. We can also see a slight improvement in NeuroQuery performance. This is due to the fact that NeuroConText and NeuroQuery models are optimized for processing long texts. In contrast, Text2Brain performs better with short descriptions, as it is not designed to handle long text effectively. The extended NeuroVault descriptions are open access in NeuroConText repository.

**Table 1.**
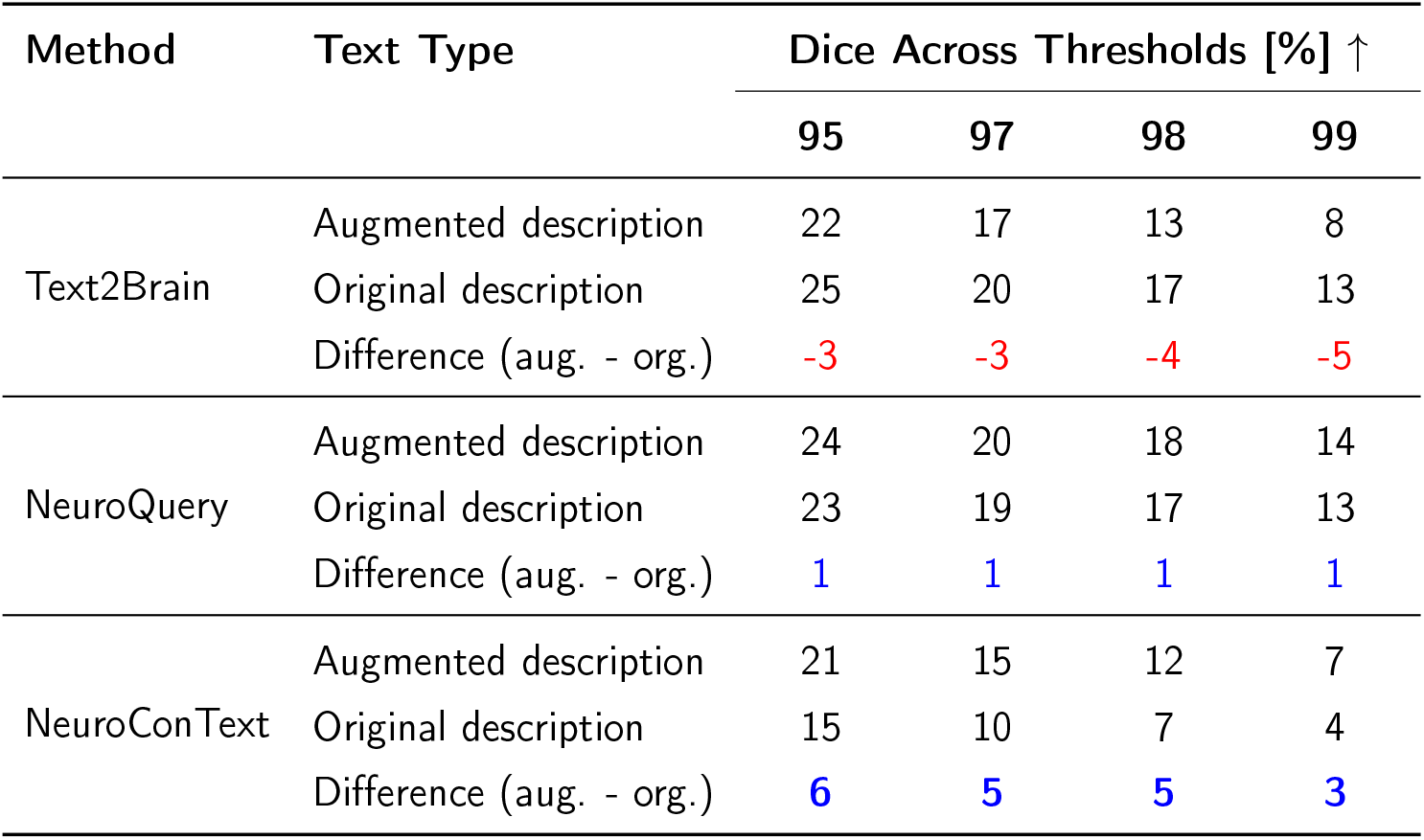
Impact of LLM-based text augmentation on reconstructing brain map from short-text input: LLM-based text augmentation improves the reconstruction performance of NeuroConText and NeuroQuery when applied to NeuroVault short labels. This confirms that these methods benefit from longer, semantically richer input text. In contrast, Text2Brain, which is optimized for short text input, performs better with the original (non-augmented) labels. This table shows the Dice scores for NeuroConText, NeuroQuery, and Text2Brain over different brain map thresholds for reconstructing NeuroVault maps from their labels. For each method, we compare the results obtained with the original short-text descriptions (org.) and the augmented descriptions (aug.).

In addition to the quantitative results, we performed a qualitative analysis of the brain maps reconstructed by NeuroConText for NeuroVault descriptions before and after augmentation, as shown in Fig.6. In this figure, we have considered three cases: samples where NeuroConText achieved the highest Dice scores, samples with median scores, and samples with the lowest scores. The comparison between long and short inputs in the best and median cases shows that reconstructions from augmented descriptions provide more focal activations, with notable overlap between maps generated by NeuroConText and the corresponding NeuroVault statistical maps. We also note that NeuroConText performs poorly on contrasts such as *0-back vs 2-back* and *evaluation of food*. This poor performance is mainly due to limitations in the labeling and missing information. For instance, in the case of *evaluation of food*, the label lacks the term “image”, e.g. *evaluation of food image*, which would imply the involvement of the visual cortex. Similarly, for *0-back vs 2-back*, is the opposite of a working memory highlighting contrast, and as such may like salient structures. As a result, NeuroConText fails to capture the upper peaks of this image.

**Figure 6.**
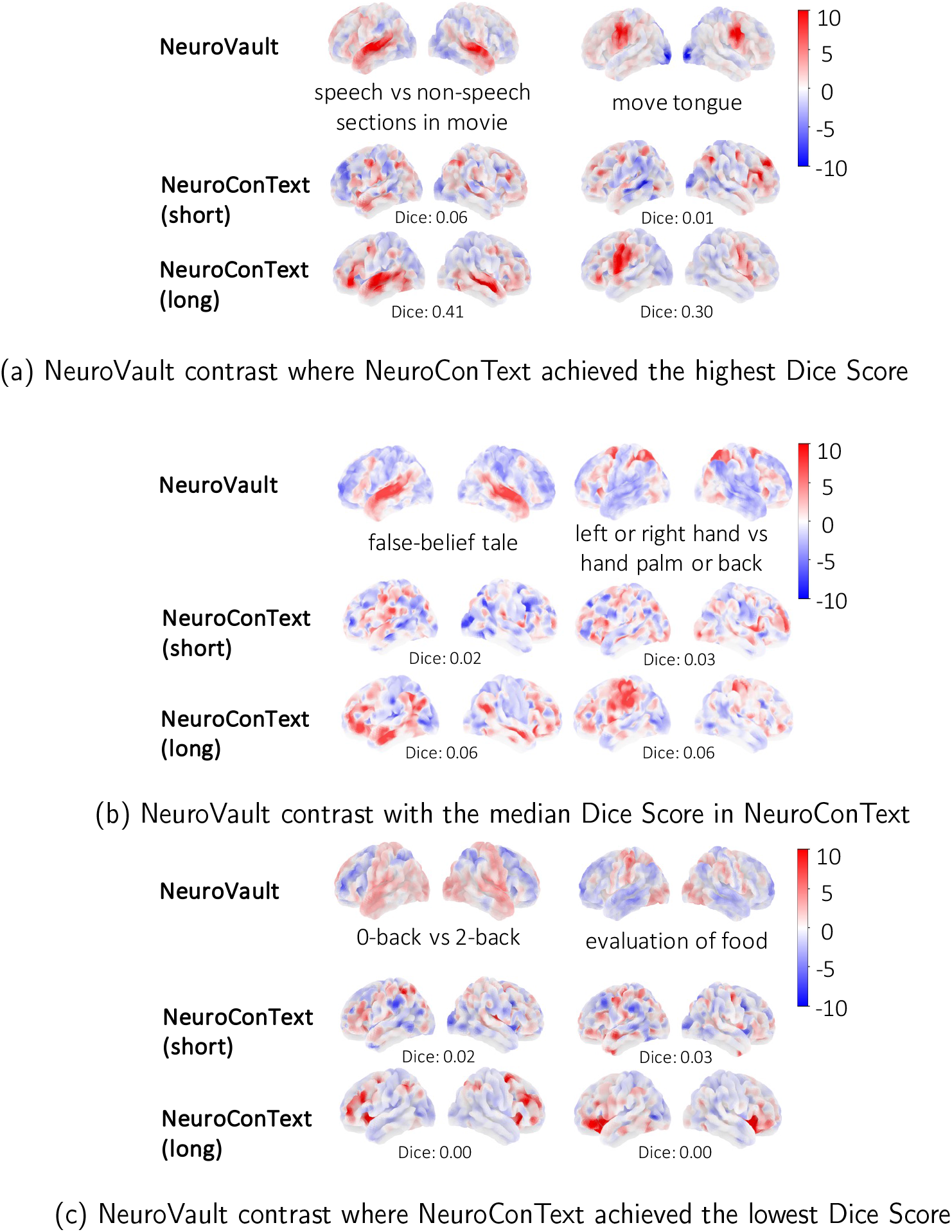
Qualitative comparison of maps reconstructed from NeuroVault descriptions: This figure presents a qualitative comparison of reconstructed brain maps for three cases where NeuroConText achieved the (a) highest, (b) median, and (c) lowest Dice scores. Each row shows the original NeuroVault statistical map and the reconstructed maps using NeuroConText with short and augmented text descriptions. In the best and median cases, reconstructions from augmented descriptions show improved localization and overlap with the NeuroVault maps. In the worst-performing cases, low-quality labeling and missing key terms contribute to poor alignment. Specifically, the *evaluation of food* label omits the word “image” leading to missing visual cortex activations, and the *0-back vs 2-back* is not expected to outline clear positive effects. NeuroConText produces more focal reconstructions when using augmented (long) descriptions, particularly in high and median performing cases.

#### 4.4.2 LLM-based text augmentation enhances NeuroConText retrieval performance on short-text

We studied the generalization of NeuroConText on retrieval tasks for short input text.^5^ The results show that NeuroConText initially does not generalize well for the short text inputs of NeuroVault descriptions. To enhance NeuroConText’s generalization on short texts, we applied our proposed LLM-based text augmentation module presented in Section 3.4. Augmented descriptions consistently improve the performance of NeuroConText both in NeuroConText’s latent space and the decoded image space across all metrics. Specifically, in the latent space, augmented descriptions lead to substantial improvements across all three metrics, with Recall@10 increasing by 9%, Recall@100 by 16%, and Mix&Match by 11% compared to the original short descriptions. In contrast, NeuroQuery and Text2Brain do not show significant improvements. In particular, Text2Brain struggles in retrieval performance on the augmented descriptions due to its limited compatibility with analyzing long text.

### 4.5 CBMA Tools Infer Regions Beyond Reported Coordinates, Mitigating the Limitation of Sparse Coordinate Reporting

In CBMA, brain activation coordinates reported in research articles are often sparse and do not capture all activated regions. This limitation affects the accuracy of the KDE maps, which rely on these coordinates to estimate full brain activation maps as it may fail to reconstruct the complete activation patterns. As CBMA tools leverage text information from research articles, aggregating both experimental descriptions and reported coordinates, we hypothesize that they have the potential to mitigate the challenge of sparse coordinate reporting in CBMA. To evaluate this hypothesis, we extracted all brain regions discussed in each article using Mistral LLM, then mapped these regions to standard brain atlases available using Nilearn(Abraham et al., 2014). We employed an *atlas label matching* procedure, in which brain region names extracted from the articles’ texts were matched to standardized labels from brain atlases. We used six atlases available in Nilearn: AAL (Tzourio-Mazoyer et al., 2002), Harvard-Oxford (“Harvard-Oxford Cortical and Subcortical Structural Atlases”, n.d.), DiFuMo (Dadi et al., 2020), Juelich (Eickhoff et al., 2007), MSDL (Varoquaux et al., 2011), and Pauli (Pauli et al., 2018). These atlases are matched with 30.2% of the articles’ extracted regions. Using the matched atlas components, we constructed an **atlas-based reference map** by normalizing and averaging the relevant components to each article. We then compared this map to both KDE-based reconstructions and NeuroConText-generated maps to assess whether NeuroConText captures active regions absent in KDE-based maps due to sparse coordinate reporting. The complete pipeline for the atlas matching process is shown in Fig.8 of Appendix 6.2.

To evaluate our approach, we performed both qualitative and quantitative analyses. For quantitative analysis, we computed Dice scores to measure spatial overlap between the atlas-based reference and both KDE-and CBMA-derived maps. Specifically, we calculated Dice scores for the (atlas, KDE) and (atlas, CBMA) pairs across different brain map thresholds and averaged the results across all articles. For qualitative analysis, we visually compared meta-analysis tools’ reconstructions with KDE-based maps and the atlas-based reference to determine whether meta-analysis tools identified regions missing from the KDE representation but present in the atlas-based reconstruction.

Our quantitative evaluation presented in Fig. 9 of Appendix 6.2 indicates that CBMA tools outperform KDE representation, with higher Dice scores for the (atlas, CBMA) pair than the (atlas, KDE) pair. This supports our hypothesis that CBMA tools have the potential to mitigate the limitations of sparse coordinate reporting in CBMA. In addition, our qualitative analysis for the best, median and worst performing samples of NeuroConText is shown in Appendix 6.2-Fig.10.^6^ The full lists of atlas-matched regions for all articles are available in the [NeuroConText GitHub repository](https://github.com/ghayem/NeuroConText/tree/main/data/Atlass_Matching). From these results, we observe that NeuroConText’s reconstructions include several active regions in the atlas-based reference but were missing from KDE-based maps. This is particularly observed in the worst-case example, where several valid activations align with the regions discussed in the article, despite the Dice score being very low. This highlights that a low Dice score, when using the KDE map as a reference, does not necessarily indicate poor reconstruction by a meta-analysis tool. Instead, it may result from the limited number of reported coordinates in the article, which underscores the limitations of KDE-based evaluation.

## 5 Discussion

In this paper, we have presented NeuroConText, a predictive brain meta-analysis tool for text-to-brain retrieval and reconstruction of brain activation maps from text, which also supports second-level CBMA by providing activation maps associated with top-K retrieved studies. It leverages a multi-objective loss using contrastive and MSE losses for retrieval and reconstruction, along with an ensemble architecture that simultaneously encodes text and brain image data in a shared latent space and decodes brain images from text latent representation. By using an advanced Large Language Model (LLM), NeuroConText captures the semantic content of entire neuroscientific articles for contextual brain map generation. NeuroConText employs LLM-based text augmentation to enhance its performance for short text lengths during inference. Our experimental results show that NeuroConText is capable of inferring additional semantic information from the text to improve the reconstructed brain map, which establishes that it effectively targets the issue that reported coordinates are often sparse and do not entirely cover all regions reported in the articles’ tables.

### Practical usage of NeuroConText for brain meta-analysis

In practice, our main proposition for neuroimaging meta-analysis is a workflow based on the best-performing configuration of NeuroConText. For a given textual query (e.g., an article, a task description, contrast, or cognitive construct), we recommend using NeuroConText to (i) retrieve the most semantically related studies, (ii) predict a corresponding consensus activation map that summarizes the spatial pattern most consistent across the literature for that query, and (iii) offer the possibility of performing second-level CBMA statistical analysis on the retrieved empirical maps. In our experiments, this workflow yields clearly improved retrieval performance compared with existing predictive brain meta-analysis baselines (e.g., NeuroQuery and Text2Brain), while reconstruction performance is on par or better than these baselines. We therefore view this NeuroConText-based retrieval–reconstruction pipeline, potentially followed by second-level CBMA on the retrieved maps, as a practical setting for users who wish to perform literature-based exploratory meta-analysis.

### Retrieval Analysis of NeuroConText

We highlight several insights derived from the quantitative and qualitative analyses of NeuroConText’s performance in the retrieval task. As shown in Fig.3, the quantitative evaluation demonstrates that NeuroConText consistently outperforms existing baselines across all text lengths and retrieval metrics. This superior performance stems from NeuroConText’s use of contrastive learning, which sets it apart from previous meta-analysis tools. Contrastive learning aligns text and the location of brain activation peaks in a shared latent space. This ensures that semantically similar texts are positioned close to their corresponding brain patterns, and that unrelated pairs are separated. This allows the model to identify the correct match between a textual description and its associated brain activity while filtering out incorrect associations. By leveraging LLMs, NeuroConText further enhances this process, as their rich, context-aware embeddings effectively handle long-text inputs and capture detailed semantics, which leads to more accurate mappings between text and brain patterns. The positive impact of the contrastive technique on retrieval performance is demonstrated in Table 2 of Supplementary Material F, where we compare it with RidgeCV and MLP models that rely solely on MSE loss. Furthermore, our results indicate that utilizing the full-body text of articles significantly enhances retrieval outcomes, which underscores the importance of leveraging complete textual information within articles.

Table 4 of Supplementary Material F illustrates the status quo regarding advanced semantic extraction. It may change in the future with the advent of novel LLMs. By using larger and more sophisticated language models, NeuroConText achieves improved semantic analysis, which in turn boosts its retrieval capabilities. This finding demonstrates the critical role of deep semantic analysis in improving the accuracy of retrieved brain activation maps.

### Reconstruction analysis of NeuroConText

In this paper, we provided both quantitative and qualitative evaluations to assess the performance of NeuroConText in reconstructing brain maps from input queries, resulting in several findings. Fig.4 quantitatively demonstrates that NeuroConText’s performance in reconstructing full-body text from left-out articles is on par with the baselines. Qualitative analyses in Fig.5 provide typical illustrations of NeuroContext behavior in view of the brain coordinates and regions reported in each article. We note that, when interpreting results from NeuroConText, it is important to distinguish between the model’s predictions and primary neuroimaging data. Given an article’s text, NeuroConText predicts the activation pattern that is most compatible with the corpus of paired text–coordinate data. This constitutes a predictive form of inference at the population level. These predicted maps are not treated as new empirical observations; they are model outputs that summarize regularities learned across studies and, in practice, can be used to generate hypotheses. They help highlight candidate regions and suggest sets of semantically related studies. Users are encouraged to interpret these predictions alongside the empirical activation maps of the retrieved studies, on which second-level statistical inference can be performed (see Appendix 6.1). We also note that NeuroConText is trained on thousands of studies containing both textual descriptions and reported coordinates, so it learns from aggregate text–activation associations rather than from the interpretation of any single article. This cross-study training emphasizes stable language–activation regularities across the literature and reduces the impact of speculative claims in individual reports. The resulting predicted maps, therefore, reflect spatial patterns that are robust across related studies whose validity can be assessed by comparison with retrieved empirical maps.

### LLM-based text augmentation impact

To accommodate short input queries typical in practical applications, we adapted NeuroConText using an LLM-based text augmentation technique. Our experiments assessed the impact of text augmentation on NeuroConText’s performance with short NeuroVault descriptions. Augmenting short texts to mimic full-length articles considerably improved the model’s reconstruction performance as shown in Table 1. We also observed that it enhanced retrieval accuracy for NeuroConText by up to 9% in Recall@10.^7^ In Fig. 6, text augmentation enhances the reconstruction task in the best- and median-performing cases, resulting in more focal activations with a noticeable overlap with the NeuroVault image. Additionally, Fig. 6(c) illustrates the least successful cases for short texts, emphasizing the challenge NeuroConText encounters when processing tasks with diffuse activations or incomplete descriptions. We note that LLMs may introduce hallucinated content during such expansions, and in our current work we do not attempt to control or filter this risk in an automated way. Instead, we assessed its impact empirically by comparing NeuroConText with and without augmentation, both at inference and on the downstream task. LLM-based augmentation consistently improves retrieval and reconstruction performance across recall@*K* and Dice Score metrics respectively, and NeuroConText with augmentation performs on par or better than the existing baselines.

Regarding the quality and potential contamination of LLM-generated content, the augmented texts and associated metadata used in our experiments are publicly available in the NeuroConText GitHub repository, and we manually reviewed them, as far as our expertise allowed, to check for correctness. In practice, we view it as the user’s responsibility to apply any additional filtering or validation appropriate to their specific use case. Moreover, developing more controlled or constrained augmentation strategies (*e*.*g*., domain-specific LLMs or dedicated filtering techniques) is an interesting direction for future work to further reduce the risk of hallucination.

### CBMA tools can infer brain activation beyond reported coordinates

CBMA methods rely on reported activation coordinates, which are often sparse and may not fully capture the underlying brain regions discussed in the articles. Many neuroscience studies report only a small number of activation peaks, which do not reflect the true spatial extent of observed activations. As a result, reconstructing full-brain activation patterns from such sparse coordinate reports and textual descriptions is intrinsically a difficult problem, and relatively modest Dice values are to be expected even for models that capture meaningful structure.

We have shown that meta-analysis tools that leverage text information and aggregate reported coordinates across thousands of articles can infer meaningful brain activations beyond the reported coordinates. To support this, in Appendix 6.2 (Fig. 8), we constructed an alternative, atlas-based reference for each article by extracting brain region names from the text using an LLM and matching them to standard atlas components. This reference includes regions discussed in the text but often missing from the reported coordinates. Our qualitative and quantitative analyses in Fig. 9 and Fig. 10 show that, in some cases, compared with the KDE map, the meta-analysis tools’ reconstructions align more closely with the atlas-based maps, capturing additional relevant regions absent in KDE-based representations. This suggests that NeuroConText is able to recover meaningful activations beyond those explicitly reported as coordinates, and that moderate Dice scores with respect to KDE maps may, in part, reflect the incompleteness and sparsity of the reported coordinate maps rather than the inadequacy of the model.

**Figure 7.**
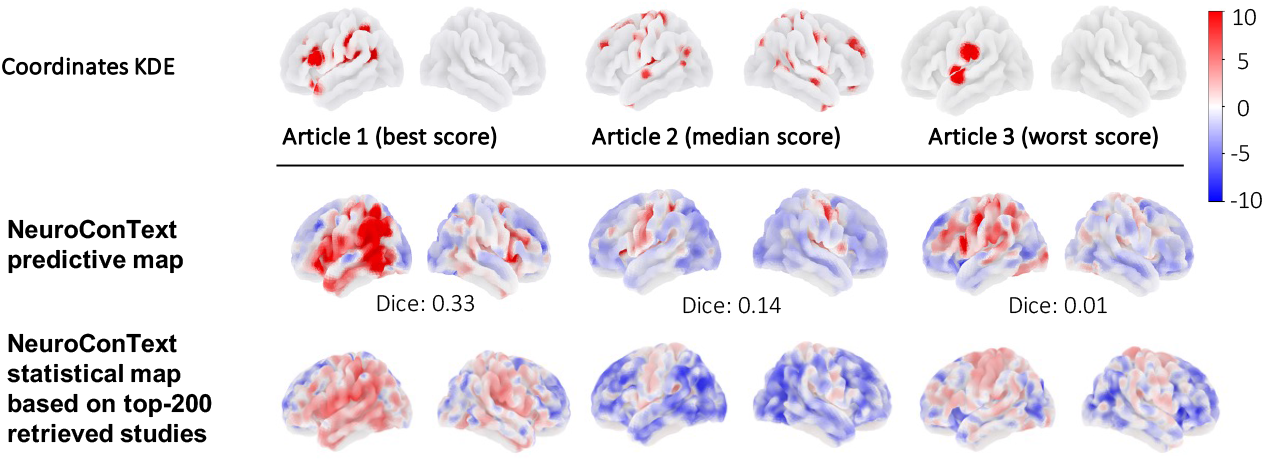
Predictive maps vs. retrieval-based second-level statistical maps.

**Figure 8.**
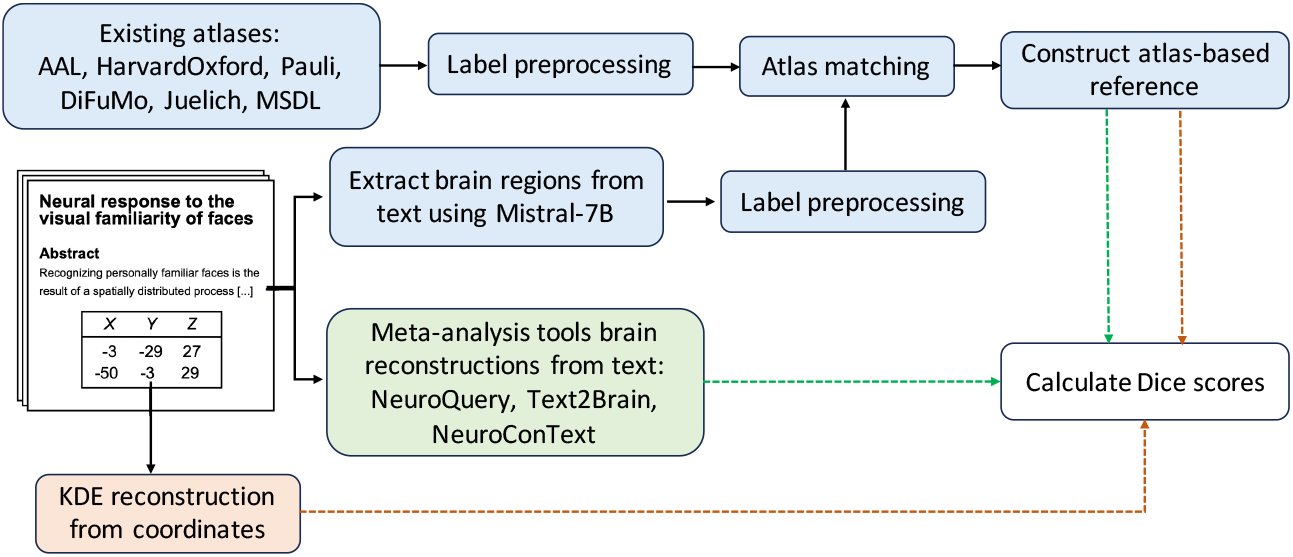
Atlas-based reference map framework to assess the reconstruction performance.

**Figure 9.**
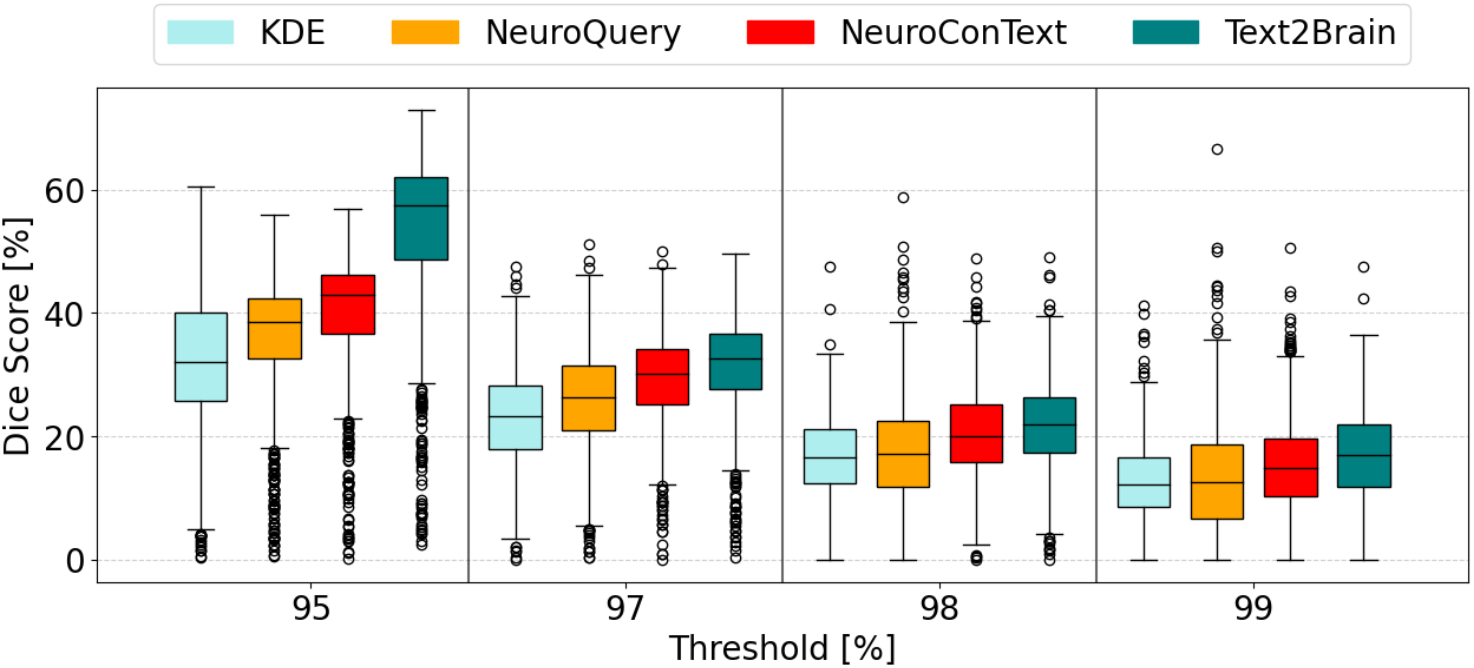
Dice similarity between atlas-based reference maps and reconstructed brain maps: Dice scores are computed between atlas-based reference maps (constructed by matching article-extracted region names to atlas components) and reconstructed brain maps generated by KDE, NeuroQuery, Text2Brain, and NeuroConText. Scores are shown across different threshold levels (95–99%) applied to the reconstructed maps. Meta-analysis tools consistently yield higher Dice scores than KDE, especially at lower thresholds. This indicates meta-analysis tools provide better overlap with the region-based reference.

**Figure 10.**
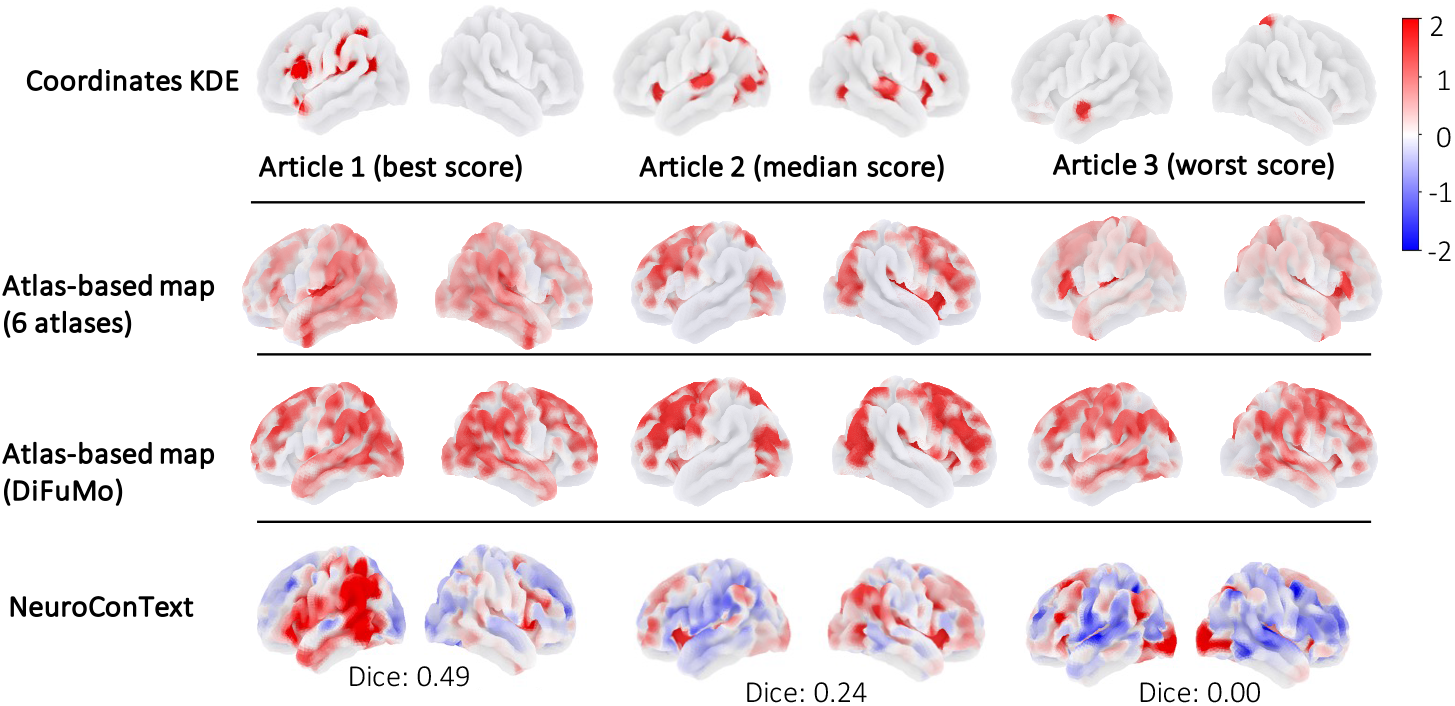
Qualitative analysis to asses NeuroConText’s ability to mitigate the sparse coordinate limitation of CBMA: Visualization of reconstructed brain maps for three representative articles with the best, median, and worst Dice scores. The first row shows KDE-based reconstructions, the second row presents the atlas-based reference maps averaged over 6 different atlases, the third row presents the atlas-based reference maps based on only DiFuMo atlas, and the last row displays NeuroConText-generated maps. The comparison highlights that NeuroConText captures broader activation patterns compared to KDE, aligning more closely with the atlas-based reference.

Overall, our results show that Dice scores computed with KDE-based coordinate maps must be interpreted with caution, as they conflate model performance with the sparsity and incompleteness of reported peaks. These findings highlight that Dice score using KDE representations of reported coordinates as reference maps provides a limited metric for evaluating CBMA tools, especially when low scores result from the inherent sparsity of reported coordinates rather than poor reconstruction. Meanwhile, comparisons with atlas-based maps can complement the analysis of reconstructed map quality and provide a more informative evaluation of CBMA tools.

### Averaged chunks, larger language models, and larger DiFuMo dictionaries boost NeuroConText

Our ablation study ^8^ demonstrated that for NeuroConText, averaging chunks of text from articles provides the best text representation, and outperforms other methods like random selection and Borda Count voting. Additionally, we showed that NeuroConText outperformed standard regression-based models (RidgeCV and non-linear models), which highlights the impact of contrastive learning to capture text-brain multimodal data relationships. Further tests showed that larger language models and dictionary sizes enhance performance, suggesting that increased model complexity and richer feature representations are beneficial for accurately mapping neuroscience texts to brain activity.

### Limitations of NeuroConText

NeuroConText faces several limitations and challenges. 1) *Text augmentation risks:* While text augmentation using LLM can enhance short-text inputs, it may introduce inaccuracies or hallucinations that require domain expert verification. This reliance on expert oversight can slow the pipeline and increase the effort needed to maintain text data integrity. 2) *Atlas-dependent evaluation for new inferred activations:* Another limitation of our study is the reliance on atlas-based label matching with articles’ extracted brain regions. We applied preprocessing steps to harmonize the atlas labels and the brain regions extracted from the articles. This approach allowed us to match 30.2% of the regions to the atlas components. However, we observed that many of the least frequently matched labels were inconsistently formatted or contained abbreviations.^9^ This highlights the importance of label quality and the need for further curation and standardization of atlas labels to improve matching coverage. 3) *Within-article experimental heterogeneity:* Individual publications often report multiple experiments, contrasts, or subsamples within a single article. In the present work, NeuroConText is evaluated at the article level, linking the full text to aggregated coordinates rather than explicitly disentangling per-table mappings. A more fine-grained analysis would require associating specific text segments with individual coordinate tables, generating separate KDE maps for each, and assessing reconstruction reliability at the within-article level. While Fig. 5 partially illustrate how NeuroConText behaves in such cases, a systematic case-study analysis of multi-experiment articles is left for future work.

## Data and Code Availability

All code and data used in this study are publicly available at: https://github.com/ghayem/NeuroConText.

## Author Contributions

**Fateme Ghayem**: Conceptualization, methodology, software, formal analysis, writing original draft, visualization.

**Raphaël Meudec**: Methodology, data preparation, software, validation, review and editing.

**Jérôme Dockès**: Data preparation, software, review and editing.

**Bertrand Thirion**: Supervision, conceptualization, funding acquisition, review and editing.

**Demian Wassermann**: Supervision, conceptualization, review and editing.

## Supplementary Material

Supplementary material related to this article is available at: https://drive.google.com/file/d/17IJ7Jn9cHXbMiBEzCnTepDcleeXHpRN-/view.

## Appendix

### 6.1 NeuroConText’s predictive map v.s. CBMA statistical map

We compare NeuroConText predictive maps with the traditional CBMA by computing a second-level statistical map: we form two study sets (top-200 retrieved vs. non-retrieved) and compute a voxel-wise second-level contrast on the KDE maps to highlight regions enriched in retrieved studies. Fig. 7 shows the target KDE map, the NeuroConText prediction, and the second-level contrast for best, median, and worst Dice scores cases. While the two often overlap, NeuroConText’s predictive maps tend to be more focal.

### 6.2 Pipeline to assess sparse coordinate mitigation in meta-analysis tools

Fig. 8 shows our pipeline to assess sparse coordinate reporting impact on Dice scores to evaluate meta-analysis ability to mitigate this limitation. Fig. 9 and Fig. 10 show quantitative and qualitative results.

## Footnotes

1 The detailed NeuroConText algorithm and the theoretical proof of convergence are provided in Supplementary Material A.

2 Refer to Supplementary Material B for a visualization of the distribution shift between short and long texts.

3 See Supplementary Material C for the prompt we used to augment NeuroVault contrast descriptions.

4 Refer to Supplementary Material B for the visualization of this distribution shift.

5 Refer to Supplementary Material D for the retrieval results.

6 The lists of articles’ regions and the matched atlas labels for the best, median, and worst scores are provided in Tables 9 to 11 of Supplementary Material G.

7 Refer to Supplementary Material D for the retrieval results.

8 All ablation results are presented in Section F of the Supplementary Material. Specifically: Table 2 compares NeuroConText with regression-based models; Table 3 compares different text aggregation methods; Table 4 evaluates the effect of LLM and DiFuMo dictionary size.

9 See Table 8 of Supplementary Material G for the list of 30 least frequent matched labels with their corresponding atlases.

