## Supplementary Material for "NeuroConText: Contrastive Learning for Neuroscience Meta-Analysis with Rich Text Representation"

December 15, 2025

#### Supplementary Material

##### A NeuroConText algorithm and its convergence proof

Proposition 1 establishes the convergence of NeuroConText algorithm, followed with the proof. We note that to set the step size $\mu$, we monitor the convergence of our approach and continuously track the decrease in InfoNCE and MSE losses, assess model performance on a validation set, and examine the norms of the gradients and the stability of parameter updates. This ensures that each optimization step contributes effectively to the overall training objective.

**Proposition 1** (Monotonic decrease of the total loss). *Consider the objective functions* $\mathcal{L}_{\mathrm{InfoNCE}}(\theta_{text}, \theta_{img})$, $\mathcal{L}_{\mathrm{MSE}}(\theta_{text}, \theta_{dec})$, *and define*

$$\mathcal{L}\big(\theta_{text}, \theta_{img}, \theta_{dec}\big) \;=\; \mathcal{L}_{\mathrm{InfoNCE}}\big(\theta_{text}, \theta_{img}\big) \;+\; \mathcal{L}_{\mathrm{MSE}}\big(\theta_{text}, \theta_{dec}\big).$$

*Assume* $\mathcal{L}_{\mathrm{InfoNCE}}$ *is* $L_1$*-smooth in* $(\theta_{text}, \theta_{img})$*, and* $\mathcal{L}_{\mathrm{MSE}}$ *is* $L_2$*-smooth in* $(\theta_{text}, \theta_{dec})$*. Then, for a sufficiently small step size* $\mu$*, the update scheme in* ***Algorithm 1*** *produces a nonincreasing sequence*

$\{\mathcal{L}(\theta_{text}^{(k)}, \theta_{img}^{(k)}, \theta_{dec}^{(k)})\}$*, which is therefore convergent to a local minimum since* $\mathcal{L}$ *is non-negative and thus lower-bounded.*

**Algorithm 1:** Alternating Minimization of $\mathcal{L}_{\text{InfoNCE}}(\theta_{\text{text}}, \theta_{\text{img}}) + \mathcal{L}_{\text{MSE}}(\theta_{\text{text}}, \theta_{\text{dec}})$

**Data:** Training data $(\mathbf{X}, \mathbf{Y})$, step size $\mu > 0$, initial parameters $\theta_{\text{text}}^{(0)}, \theta_{\text{img}}^{(0)}, \theta_{\text{dec}}^{(0)}$

**Result:** Final parameters $\theta_{\text{text}}^{(k)}, \theta_{\text{img}}^{(k)}, \theta_{\text{dec}}^{(k)}$

**while** *stopping criterion not met* **do**

/* Step 1: Intermediate update of $\theta_{\text{text}}$ w.r.t $\mathcal{L}_{\text{InfoNCE}}$ */

$\tilde{\theta}_{\text{text}}^{(k+1)} \leftarrow \theta_{\text{text}}^{(k)} - \mu \nabla_{\theta_{\text{text}}} \mathcal{L}_{\text{InfoNCE}} \left( \theta_{\text{text}}^{(k)}, \theta_{\text{img}}^{(k)} \right)$;

/* Step 2: Update of $\theta_{\text{img}}$ w.r.t $\mathcal{L}_{\text{InfoNCE}}$ */

$\theta_{\text{img}}^{(k+1)} \leftarrow \theta_{\text{img}}^{(k)} - \mu \nabla_{\theta_{\text{img}}} \mathcal{L}_{\text{InfoNCE}} \left( \theta_{\text{text}}^{(k)}, \theta_{\text{img}}^{(k)} \right)$;

/* Step 3: Final update of $\theta_{\text{text}}$ w.r.t $\mathcal{L}_{\text{MSE}}$ */

$\theta_{\text{text}}^{(k+1)} \leftarrow \tilde{\theta}_{\text{text}}^{(k+1)} - \mu \nabla_{\theta_{\text{text}}} \mathcal{L}_{\text{MSE}} \left( \tilde{\theta}_{\text{text}}^{(k+1)}, \theta_{\text{dec}}^{(k)} \right)$;

/* Step 4: Update of $\theta_{\text{dec}}$ w.r.t $\mathcal{L}_{\text{MSE}}$ */

$\theta_{\text{dec}}^{(k+1)} \leftarrow \theta_{\text{dec}}^{(k)} - \mu \nabla_{\theta_{\text{dec}}} \mathcal{L}_{\text{MSE}} \left( \theta_{\text{text}}^{(k+1)}, \theta_{\text{dec}}^{(k)} \right)$;

$k \leftarrow k + 1$;

/* Check stopping criterion */

**end**

***Proof***. Let's define $\theta_1 := \theta_{\text{text}}$, $\theta_2 := \theta_{\text{img}}$, $\theta_3 := \theta_{\text{dec}}$ and two sub-losses $f(\theta_1, \theta_2) = \mathcal{L}_{\text{InfoNCE}}(\theta_1, \theta_2)$ and $g(\theta_1, \theta_3) = \mathcal{L}_{\text{MSE}}(\theta_1, \theta_3)$. Then, the total objective becomes:

$$\mathcal{L}(\theta_1, \theta_2, \theta_3) = f(\theta_1, \theta_2) + g(\theta_1, \theta_3).$$

We divide the proof into two parts: (1) showing that the $\theta_1, \theta_2$ updates (Steps 1–2 in Algorithm 1) reduce $f$; and (2) showing that the $\theta_1, \theta_3$ updates (Steps 3–4 in Algorithm 1) reduce $g$. We then argue that these reductions are not undone by the intervening updates, provided $\mu$ is sufficiently small.

**1. Steps 1–2 decrease $f(\theta_1, \theta_2)$:**

**Step 1 (gradient descent on $\theta_1$ wrt $f$)**:

$$\tilde{\theta}_1^{(k+1)} = \theta_1^{(k)} - \mu \nabla_{\theta_1} f\big(\theta_1^{(k)}, \theta_2^{(k)}\big).$$

Under $L_f$-smoothness of $f$ and for $\mu \le 1/L_f$, standard gradient-descent theory Bertsekas, 1997; Nesterov et al., 2018 guarantees

$$f\big(\tilde{\theta}_1^{(k+1)}, \theta_2^{(k)}\big) \;\le\; f\big(\theta_1^{(k)}, \theta_2^{(k)}\big) \;-\; \Delta_{f,1}^{(k)},$$

where $\Delta_{f,1}^{(k)} > 0$ depends on $\mu$ and $\|\nabla_{\theta_1} f(\cdot)\|^2$.

**Step 2 (gradient descent on $\theta_2$ wrt $f$)**:

$$\theta_2^{(k+1)} \;=\; \theta_2^{(k)} \;-\; \mu\,\nabla_{\theta_2} f\big(\theta_1^{(k)}, \theta_2^{(k)}\big).$$

By $L_f$-smoothness in $\theta_2$, we get

$$f\big(\theta_1^{(k)}, \theta_2^{(k+1)}\big) \;\le\; f\big(\theta_1^{(k)}, \theta_2^{(k)}\big) \;-\; \Delta_{f,2}^{(k)},$$

for some positive $\Delta_{f,2}^{(k)}$. Combining these sub-steps yields

$$f\big(\tilde{\theta}_1^{(k+1)}, \theta_2^{(k+1)}\big) \;\le\; f\big(\theta_1^{(k)}, \theta_2^{(k)}\big) \;-\; \Delta_f^{(k)}, \quad \Delta_f^{(k)} = \Delta_{f,1}^{(k)} + \Delta_{f,2}^{(k)} > 0.$$

Hence, after Steps 1–2, $f$ is strictly decreased from its value at the start of iteration $k$.

2. Steps 3–4 Decrease $\boldsymbol{g(\theta_1, \theta_3)}$.

**Step 3 (gradient descent on $\theta_1$ wrt $g$)**:

$$\theta_1^{(k+1)} \;=\; \tilde{\theta}_1^{(k+1)} \;-\; \mu\,\nabla_{\theta_1} g\big(\tilde{\theta}_1^{(k+1)}, \theta_3^{(k)}\big).$$

If $g$ is $L_g$-smooth in $\theta_1$ and $\mu \le 1/L_g$, then

$$g\big(\theta_1^{(k+1)}, \theta_3^{(k)}\big) \;\le\; g\big(\tilde{\theta}_1^{(k+1)}, \theta_3^{(k)}\big) \;-\; \Delta_{g,1}^{(k)}.$$

**Step 4 (gradient descent on $\theta_3$ wrt $g$)**:

$$\theta_3^{(k+1)} \;=\; \theta_3^{(k)} \;-\; \mu\,\nabla_{\theta_3} g\big(\theta_1^{(k+1)}, \theta_3^{(k)}\big).$$

By the same $L_g$-smoothness argument,

$$g\big(\theta_1^{(k+1)}, \theta_3^{(k+1)}\big) \;\le\; g\big(\theta_1^{(k+1)}, \theta_3^{(k)}\big) \;-\; \Delta_{g,2}^{(k)},$$

so combining both sub-steps gives

$$g\big(\theta_1^{(k+1)}, \theta_3^{(k+1)}\big) \;\le\; g\big(\tilde{\theta}_1^{(k+1)}, \theta_3^{(k)}\big) \;-\; \Delta_g^{(k)}, \quad \Delta_g^{(k)} = \Delta_{g,1}^{(k)} + \Delta_{g,2}^{(k)} > 0.$$

Thus, after Steps 3–4, $g$ is strictly decreased from its value at the end of Step 2.

##### 3. Ensuring the second update of $\theta_1$ (Step 3) cannot undo the decrease in $f$.

A critical concern is that $\theta_1$ is changed *twice* in the same iteration: once w.r.t. $f$ (Step 1), once w.r.t. $g$ (Step 3). In the following, we prove that by choosing $\mu$ sufficiently small, the second update does not cause $f$ to increase again. In particular, a common sufficient condition is

$$0 < \mu \;<\; \frac{1}{L_f + L_g} \quad \text{or} \quad 0 < \mu \;<\; \frac{1}{2 \times \max(L_f,\, L_g)}.$$

##### 3.1 First-order vs. second-order effects.

1. *Direct update on $f$* (Step 1) is a *first-order* improvement. By a first-order Taylor expansion Rudin et al., 1964:
$$f(\theta_1 - \mu\,\nabla f(\theta_1)) \;\approx\; f(\theta_1) \;-\; \mu\,\|\nabla f(\theta_1)\|^2,$$
indicating a change proportional to $\mu\,\|\nabla f\|$.

2. *Change in $f$ due to a step on $g$* (Step 3) is a *second-order* Lipschitz effect. If $\Delta\theta_1 = -\mu\,\nabla g$, then by $L_f$-smoothness:
$$\big|\Delta f\big| \;=\; \big|f(\theta_1 + \Delta\theta_1) - f(\theta_1)\big| \;\le\; \tfrac{L_f}{2}\,\|\Delta\theta_1\|^2 \;=\; \mathcal{O}(\mu^2).$$

Hence, the first-order decrease from Step 1 (of order $\mu$) dominates the second-order bump (of order $\mu^2$) for sufficiently small $\mu$. In practice, one often takes $\mu < 1/(L_f + L_g)$ so that $\mu^2 \ll \mu$. Similarly, Steps 1–2 cannot undo progress on $g$ from Steps 3–4.

**4. Conclusion: Non-increase of $\mathcal{L}$ in one full iteration.** Therefore, if $\mu$ is small enough,

$$f\big(\theta_1^{(k+1)}, \theta_2^{(k+1)}\big) + g\big(\theta_1^{(k+1)}, \theta_3^{(k+1)}\big) \leq f\big(\theta_1^{(k)}, \theta_2^{(k)}\big) + g\big(\theta_1^{(k)}, \theta_3^{(k)}\big).$$

As a result,

$$\mathcal{L}\big(\theta_1^{(k+1)}, \theta_2^{(k+1)}, \theta_3^{(k+1)}\big) \leq \mathcal{L}\big(\theta_1^{(k)}, \theta_2^{(k)}, \theta_3^{(k)}\big).$$

Since $\mathcal{L}$ is lower-bounded (non-negative), this non-increasing sequence converges to a finite limit. □

### B Visualizing the distribution shift between short and long text

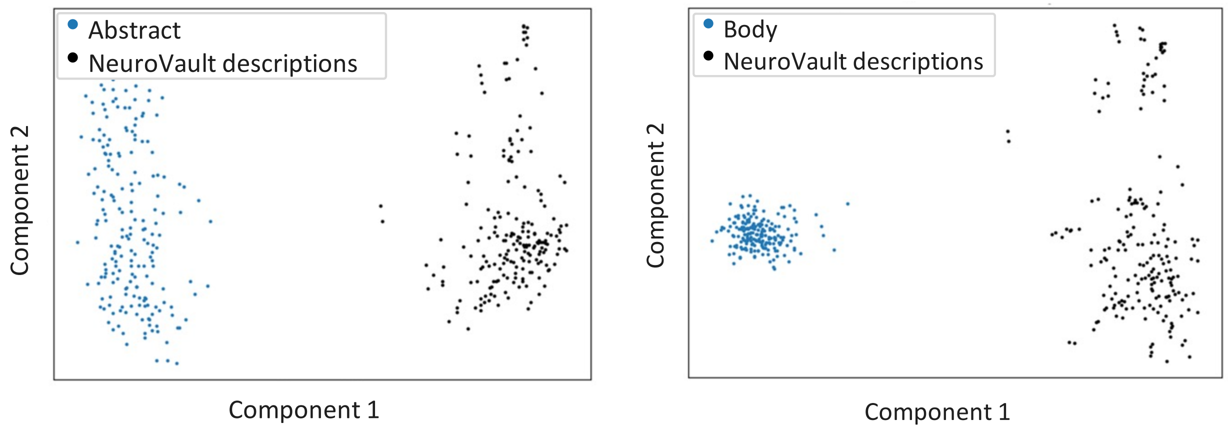


(a) PCA: NeuroVault contrasts vs abstracts

(b) PCA: NeuroVault contrasts vs body

Figure 1: PCA decomposition of LLM embeddings comparing NeuroVault contrast definitions (short text) with abstracts and full-body texts (long text). The left plot shows a clear distribution shift along the first component between NeuroVault descriptions and abstracts, while the right plot shows a more pronounced shift along both the first and second components between NeuroVault descriptions and full-body texts. This indicates a significant distribution shift between short NeuroVault descriptions and full-length articles.

#### C Prompt for augmenting NeuroVault contrast descriptions

Below is the prompt we used to extend NeuroVault contrast descriptions with GPT-4o:

```
I have 169 descriptions of the NeuroVault contrasts dataset for the brain. On
the other hand, I have trained an autoencoder to encode the brain text and
images from neuroscientific articles and to decode images from the text's
latent (encoded) representation. Now, I want to use my trained autoencoder to
encode the NeuroVault descriptions and decode (estimate) the NeuroVault brain
contrasts. However, the NeuroVault descriptions are very short, while my
autoencoder networks have been trained on lengthy neuroscientific articles.
Therefore, I want to extend the NeuroVault descriptions to resemble the body of
 an article, including sections like introduction, framework/method,
experiments, discussion, conclusion, etc., similar to an 8-page or longer
article. I will provide you with the NeuroVault descriptions, and I would like
you to extend them for me. I want the text to be as long as possible and to
include all the relevant key terms related to the description.
```

### D The impact of LLM-based text augmentation on the performance of the retrieval task

Table 1: Retrieval task on NeuroVault: the impact of NeuroConText contrastive approach and LLM-based text augmentation. Augmented descriptions consistently improve the performance of NeuroConText (both in latent and decoded forms) across all metrics—Recall@10, Recall@100, and Mix&Match—indicating that LLM-based text augmentation enhances the model's ability to retrieve relevant brain maps. For NeuroConText, augmented descriptions result in significant gains in Recall@10 (+9%), Recall@100 (+16%) and Mix&Match (+11%) compared to original descriptions. Similarly, NeuroConText (decoded) benefits from augmentation, showing improvements of +10%, +7%, and +8% across the three metrics, respectively. In contrast, NeuroQuery and Text2Brain do not show significant improvements, with Text2Brain showing slight declines across all metrics when using augmented descriptions.

| **Method** | Metric [%] ↑ / **Description Type** | **Recall@10** | **Recall@100** | **Mix&Match** |
|---|---|---|---|---|
| **NeuroConText** (in latent) | Augmented description | 13 | 71 | 64 |
| | Original description | 4 | 55 | 53 |
| | Difference (aug. - org.) | 9 | 16 | 11 |
| **NeuroConText** (decoded) | Augmented description | 20 | 70 | 66 |
| | Original description | 10 | 63 | 58 |
| | Difference (aug. - org.) | 10 | 7 | 8 |
| **NeuroQuery** | Augmented description | 20 | 70 | 65 |
| | Original description | 20 | 67 | 63 |
| | Difference (aug. - org.) | 0 | 3 | 2 |
| **Text2Brain** | Augmented description | 12 | 61 | 58 |
| | Original description | 13 | 67 | 62 |
| | Difference (aug. - org.) | -1 | -6 | -4 |

### E Prompt used for article summarization

We used a customized prompt to extract essential information about brain regions and associated cognitive processes to generate summaries of neuroscientific articles. The prompt instructed the latest version of the Mistral language model to act as a neuroscientific expert, asking relevant questions about the article and providing answers. Below is the exact prompt used:

```
You are a neuroscientific expert. Your task is to write a summary of a
neuroscientific article. When presented with the article, come up with
interesting questions to ask and answer each question. Afterward, combine all
the information and write a summary.

article: {essay}
Instructions:
*Summarize: In clear and concise language, summarize the key points and themes
presented in the article.

*Write a short summary: Write a short summary in 2 sentences.

*The brain regions: Write the brain regions and any cognitive or affective
processes related to brain activation.
```

### F Validating NeuroConText Elements

#### F.1 NeuroConText vs regression-based strategies

We compared NeuroConText with regression-based methods to associate texts with brain maps. For NeuroConText, we retrieve the text and coordinates latents of the test set through NeuroConText's trained encoders. Then, we calculate retrieval metrics, Recall@K for $K \in \{10, 100\}$, and Mix&Match, on the test data. As a regression-based approach, we consider two models: RidgeCV (linear model) and a non-linear model. For the non-linear model, we implemented a feedforward neural network comprising three fully connected layers of 512 units. Non-linear transformations are introduced through hyperbolic tangent (tanh) activation functions following the first two layers. To enhance training stability, layer normalization is applied after each activation. In addition, a dropout layer with a 50% rate is incorporated after the first activation to mitigate overfitting. This model was trained over 50 epochs using the Adam optimizer (Kingma & Ba, 2014) with a learning rate of $5e^{-4}$. We train these two models on the average Mistral-7B embeddings of the full-body chunks to estimate the DiFuMo coefficients. In Table 2, we compare the association performance of the trained models on test data and compared them with those obtained from NeuroConText and the baseline models. We can see that regression-based methods fail to associate text with brain locations effectively.

Table 2: **Comparison of NeuroConText contrastive model with regression-based models for retrieval task:** NeuroConText outperforms regression-based models (RidgeCV and a non-linear regressor) in associating texts with brain maps. This table presents the retrieval performance of NeuroConText, RidgeCV (linear regression), and a non-linear model on the test set articles using full-body text. The evaluation metrics include Recall@10, Recall@100, and Mix&Match. NeuroConText achieves the highest scores across all metrics (Meudec et al., 2024).

| Method | Recall@10 [%] | Recall@100 [%] | Mix&Match [%] |
|---|---|---|---|
| NeuroConText (ours) | $\mathbf{22.6 \pm 1.4}$ | $\mathbf{57.8 \pm 1.6}$ | $\mathbf{84.2 \pm 0.9}$ |
| RidgeCV (linear model) | $14.9 \pm 0.8$ | $43.7 \pm 1.2$ | $76.5 \pm 0.4$ |
| Non-linear model | $11.4 \pm 0.4$ | $39.4 \pm 1.2$ | $74.1 \pm 0.8$ |

##### F.2 Long-Text Chunks Aggregation: Mean Aggregation Outperforms Other Strategies

Here, we assess the chunk aggregation strategy for long texts by comparing alternative aggregation methods. We compare the following methods:

- **Random chunk selection:** We selected one random chunk from each article. By choosing a single chunk at random, we avoid introducing any bias from specific parts of the article. We repeated this random selection multiple times to ensure the robustness and consistency of the results.

- **Train-set augmentation over chunks:** To expand the dataset, we construct all possible (chunk, DiFuMo) pairs for each article, where each article has multiple text chunks but is associated with a single DiFuMo. During training, we ensure that no more than one chunk from the same article appears in a batch, maintaining a diagonal structure in the similarity matrix between text and coordinates representation in NeuroConText latent space. For the test set, we explored two approaches while ensuring that the similarity matrix maintains a diagonal structure: 1) randomly selecting one chunk per article and 2) using the average of all chunks per article.

- **Voting strategy via Borda Count:** We use the Borda Count (Dwork et al., 2001) approach for the voting strategy, where each chunk from an article produces a ranked list of brain maps. These rankings reflect how likely each brain map is to match the content of that chunk. To make a final prediction for the article, we apply the Borda Count by combining the ranks from all chunks: brain maps that appear near the top of many chunk-level rankings receive higher overall scores. The brain map with the best overall rank is selected as the final prediction. We evaluate this strategy using recall metrics to assess how well it identifies the correct brain map.

Table 3: **Comparison of chunk aggregation strategies for long-text input in NeuroConText:** Averaging text chunks' embeddings yields the best performance in associating long neuroscientific articles with brain maps. This strategy outperforms others by producing more compact and representative embeddings. The table reports retrieval performance (Recall@10, Recall@100, and Mix&Match) for various chunk aggregation strategies in NeuroConText, including random chunk selection, train-time augmentation with different test-time strategies, and a voting scheme using Borda Count.

| Chunk Aggregation Method | Recall@10 [%] | Recall@100 [%] | Mix&Match [%] |
|---|---|---|---|
| Train: augment – Test: average chunks | $20.5 \pm 1.7$ | $53.5 \pm 1.6$ | $81.9 \pm 0.5$ |
| Train: augment – Test: random chunk | $17.6 \pm 1.2$ | $49.9 \pm 1.4$ | $79.4 \pm 0.6$ |
| Voting (Borda Count) | $18.5 \pm 0.8$ | $51.9 \pm 1.9$ | $81.4 \pm 0.5$ |
| Averaging chunks | $\mathbf{22.6 \pm 1.4}$ | $\mathbf{57.8 \pm 1.6}$ | $\mathbf{84.2 \pm 0.9}$ |

Table 3 shows a comparison the four different aggregation strategies based on evaluation metrics Recall@10, Recall@100, and Mix&Match. From this table, we conclude that averaging chunks performs the best. This is possibly due to the fact that averaging creates embeddings that are more tightly grouped within chunks of an article while maintaining a clear separation between different articles' chunks embeddings in the latent space. To explore this, in Fig. 2 we used **UMAP projections** to visualize the clustering of embeddings. The UMAP visualizations show that the embeddings of any given text form compact groups, that are clearly separated between articles. The small distances within these average embeddings highlight strong grouping, which implies that averaging leads to a good representative of each article's multiple embeddings.

#### F.3 Impact of large language model and DiFuMo size

In our final experiment, we compare different LLMs and DiFuMo sizes on the performance of our proposed method. In NeuroConText model, we change the LLM to GPT-Neo-1.3B, GPT-Neo-125M, or SciBERT, alongside two DiFuMo sizes: $k = 256$ and $k = 512$. Then we compare the impact of each on the retrieval performance through the metrics Recall@K for $K \in \{10, 100\}$, and Mix&Match. The findings are summarized in Table 4. We see that the larger language model with Mistral-7B significantly enhances performance, achieving a score more than 4% higher than SciBERT. Moreover, the results indicate that the DiFuMo with size 512 outperforms 256, improving scores by approximately 3%.

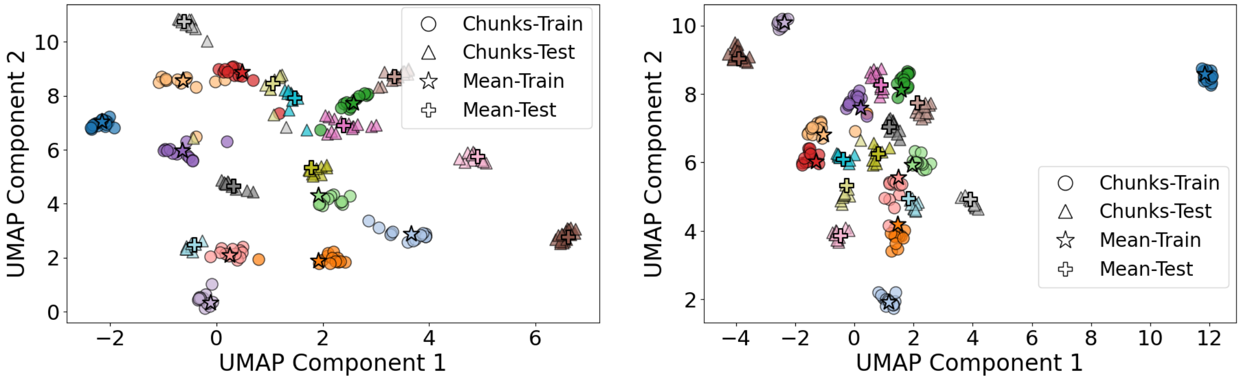


(a) Original Mistral-7B embeddings space

(b) NeuroConText text latent space

Figure 2: **UMAP visualizations of the top 10 articles with the highest number of chunks.** The left figure shows the original Mistral-7B embeddings and the right figure shows the encoded text embeddings in the shared latent space of NeuroConText model. Each color corresponds to the chunks of a given article. In both representations, chunks corresponding to the same article are closely clustered around their mean, while the means of different articles remain well-separated. This pattern supports the effectiveness of the averaging strategy, as it preserves proximity within an article's chunks to their mean while maintaining clear separation from the chunks of other articles in the latent space.

Table 4: **Impact of LLM and DiFuMo size on retrieval performance:** Using a larger language model (Mistral-7B) and a higher-resolution DiFuMo atlas (512 components) improves retrieval performance in NeuroConText. This table presents the retrieval scores for different LLMs (Mistral-7B, GPT-Neo-1.3B, GPT-Neo-125M, and SciBERT) combined with DiFuMo sizes (256 vs. 512). The evaluation metrics are Recall@10, Recall@100, and Mix&Match. Mistral-7B consistently achieves the best performance, while increasing DiFuMo resolution improves results by 3% on average (Meudec et al., 2024).

| DiFuMo Size | LLM | Recall@10 [%] | Recall@100 [%] | Mix&Match [%] |
|---|---|---|---|---|
| 256 | Mistral-7B | $\mathbf{19.8 \pm 0.9}$ | $\mathbf{51.6 \pm 1.3}$ | $\mathbf{81.0 \pm 0.6}$ |
| | GPT-Neo-1.3B | $18.1 \pm 0.6$ | $48.2 \pm 1.3$ | $79.4 \pm 0.3$ |
| | GPT-Neo-125M | $15.1 \pm 0.6$ | $42.3 \pm 1.3$ | $76.4 \pm 0.3$ |
| | SciBERT | $15.1 \pm 0.6$ | $42.8 \pm 0.9$ | $76.9 \pm 0.3$ |
| 512 | Mistral-7B | $\mathbf{22.6 \pm 1.4}$ | $\mathbf{57.8 \pm 1.6}$ | $\mathbf{84.2 \pm 0.9}$ |
| | GPT-Neo-1.3B | $21.5 \pm 1.1$ | $54.8 \pm 1.1$ | $82.7 \pm 0.5$ |
| | GPT-Neo-125M | $17.5 \pm 1.1$ | $48.2 \pm 1.5$ | $79.7 \pm 0.7$ |
| | SciBERT | $17.9 \pm 0.8$ | $50.3 \pm 1.5$ | $81.0 \pm 0.8$ |

### G Technical details of atlas label matching and coverage statistics

We applied a preprocessing and normalization procedure to improve alignment between article-extracted brain region names and atlas labels. This included standardizing label formats (Table 5) and enforcing hemisphere-specific matching. For example, "left Hippocampus" is only compared to left-sided atlas components (Table 6). Matching was performed using six atlases from Nilearn: AAL, Harvard-Oxford, DiFuMo, Juelich, MSDL, and Pauli. Note that we consider more directional restrictions when matching article region names to atlas labels; our pipeline considers that all terms (words) from the article region must appear in the atlas label after standardization. This includes directional terms like "superior", "inferior", "anterior", and "posterior". If such a term is present in the article region but missing in the atlas label, the match fails. Table 7 provides an example for matching with directional terms. With this preprocessing, we obtained 30.21% coverage of the articles' regions by the 6 atlas labels. Fig. 3 presents the distribution of matched brain region labels across different atlases, showing that the majority of matches (over 90%) originate from the DiFuMo atlas. Also, the list of 30 most frequently matched labels (all associated with DiFuMo atlas), and the 30 least frequent matched labels are shown in Figure 4 and Table 8, respectively.

Table 5: Text preprocessing rules for atlas labels and article regions.

| **Before** | **After** |
|---|---|
| LH, lh, _lh, _l | left |
| RH, rh, _rh, _r | right |
| hemi, h, hemisphere | hemisphere |
| gyri | gyrus |
| cortex | (removed) |
| lobe | (removed) |
| hemisphere | (removed) |
| Extra spaces | Removed |
| All text | Lowercased |

Table 6: Hemisphere matching between article regions and atlas components

| **Article region hemisphere** | **Assigned atlas component hemisphere.** |
|---|---|
| right | right |
| left | left |
| left and right | left, right, right & left, non |
| non | left, right, right & left, non |

Table 7: Examples of region matching with directional terms.

| Article Region | Atlas Label | Match? |
|---|---|---|
| superior frontal gyrus | superior frontal gyrus | Yes |
| superior frontal gyrus | frontal gyrus | No |
| frontal gyrus | superior frontal gyrus | Yes |
| posterior cingulate | posterior cingulate cortex | Yes |
| posterior cingulate | cingulate cortex | No |
| inferior parietal lobule | inferior parietal lobule | Yes |
| inferior parietal lobule | parietal lobule | No |

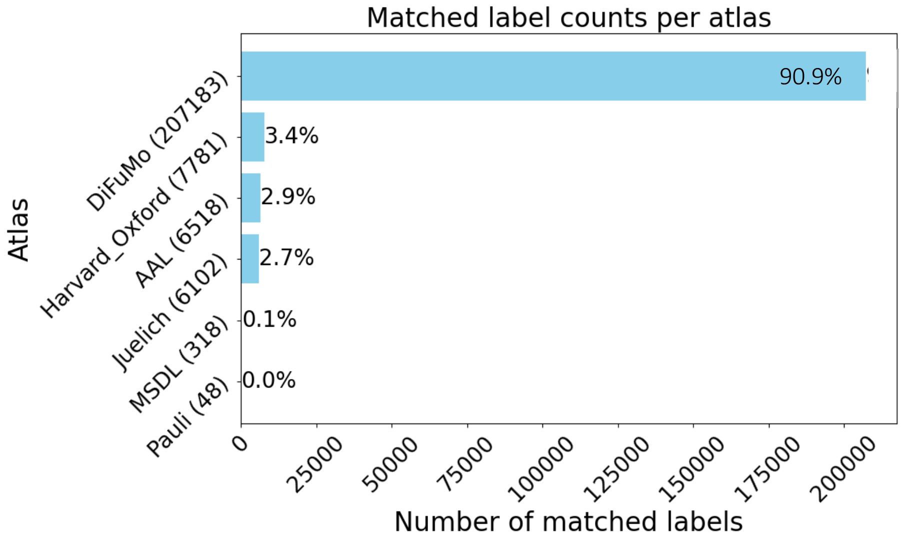


Figure 3: Number of matched brain regions per atlas across all articles.

Tables 9, 10, and 11 report the brain regions extracted from the articles using a large language model (LLM), followed by the matched atlas labels for Articles 1, 2, and 3, respectively. The matching was performed across six different atlases. The full lists of atlas-matched regions are available in the NeuroConText GitHub repository.

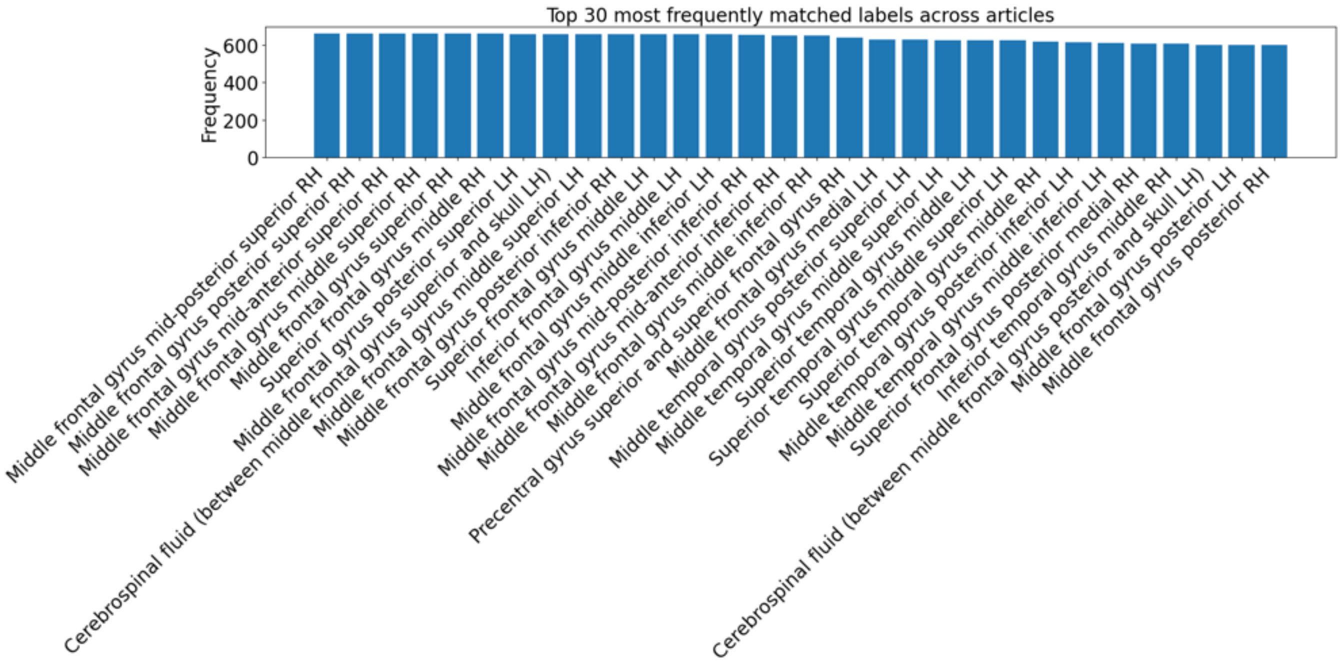


Figure 4: List of 30 most frequently matched labels.

Table 8: 30 least frequent matched labels with their corresponding atlases

| **Label** | **Count** | **Atlas** |
|---|---|---|
| Vermis_10 | 2 | AAL |
| Vermis_1_2 | 2 | AAL |
| Vermis_3 | 2 | AAL |
| Vermis_4_5 | 2 | AAL |
| Vermis_6 | 2 | AAL |
| Vermis_7 | 2 | AAL |
| Vermis_8 | 2 | AAL |
| Vermis_9 | 2 | AAL |
| Suborbital sulcus | 2 | DiFuMo |
| WM Acoustic radiation | 2 | Juelich |
| WM Uncinate fascicle | 2 | Juelich |
| Broca | 2 | MSDL |
| Med DMN | 2 | MSDL |
| STH | 2 | Pauli |
| GM Amygdala_centromedial group | 3 | Juelich |
| GM Amygdala_laterobasal group | 3 | Juelich |
| GM Amygdala_superficial group | 3 | Juelich |
| Basal | 3 | MSDL |
| Cereb | 3 | MSDL |
| Cing | 3 | MSDL |
| Front DMN | 3 | MSDL |
| Occ post | 3 | MSDL |
| Striate | 3 | MSDL |
| Sup Front S | 3 | MSDL |
| Vis | 3 | MSDL |
| Ca | 3 | Pauli |
| EXA | 3 | Pauli |
| GPe | 3 | Pauli |
| GPi | 3 | Pauli |
| HN | 3 | Pauli |

Table 9: Brain regions and atlas label matching for **Article 1 - (best Dice score)**. The full list is available in the NeuroConText GitHub repository for *PMID 30210434*.

| Source | Sample Regions |
|---|---|
| **Article regions extracted via Mistral LLM** | **Left Ventro-dorsal Stream**: Inferior parietal lobule (incl. Supramarginal gyrus), Inferior frontal gyrus (ventral premotor cortex, opercular, triangular), Precentral gyrus, Rolandic operculum, Insula.<br>**Left Frontal Cortex**: Precentral gyrus, Superior/Middle/Inferior frontal gyri (incl. opercular, triangular, orbital), Rolandic operculum.<br>**Left Parietal Cortex**: Postcentral gyrus, Superior and Inferior parietal lobules (incl. Supramarginal and Angular gyri).<br>**Left Temporal Cortex**: Superior and Middle temporal gyri, Superior temporal pole.<br>**Subcortical Regions**: Putamen, Pallidum, Thalamus.<br>**Other**: Left Insula, Left Angular Gyrus. |
| **AAL** | Pallidum_L, Pallidum_R, Putamen_L, Putamen_R, Thalamus_L, Thalamus_R |
| **Harvard-Oxford** | Angular Gyrus, Frontal Medial Cortex, Intracalcarine Cortex, Middle Temporal Gyrus (anterior division), Planum Temporale, Superior Frontal Gyrus |
| **DiFuMo (1024)** | Angular gyrus anterior LH, Frontal pole medial, Fusiform gyrus posterior RH, Lingual gyrus posterior LH, Middle frontal gyrus mid-anterior RH, Thalamus superior. *Full region mapping available in NeuroConText GitHub repository.* |
| **Juelich** | GM Parietal operculum OP1–OP4, GM Superior parietal lobule 7A, 7PC |

Table 10: Brain regions and atlas label matching for **Article 2 - (median Dice score)**. The full list is available in the NeuroConText GitHub repository for *PMID 36166644*.

| Source | Sample Regions |
|---|---|
| **Article regions extracted via Mistral LLM** | **Frontal Regions**: Frontal to central areas, Medial frontal area, Middle frontal gyrus (incl. right), Inferior frontal gyrus, Anterior insula (incl. dorsal anterior and right dorsal anterior insula), Precentral gyrus, Rolandic operculum.<br>**Parietal Regions**: Inferior and Superior parietal lobules, General parietal areas.<br>**Temporal Regions**: Superior temporal gyrus (bilateral and posterior), Superior temporal plane (STP), Planum temporale (PT), Temporo-parieto-occipital junction (TPO).<br>**Occipital Regions**: Occipital areas, Inferior occipital gyrus, Visual cortex.<br>**Cingulate Regions**: Anterior cingulate cortex (ACC), Dorsal ACC (dACC), Midcingulate cortex (MCC).<br>**Other**: Fusiform gyrus, Optic nerve, Optic chiasm, Salience network, Superior olivary nucleus, Right/Left-/Bilateral hemispheres, Contralateral and Ipsilateral attention systems. |
| **AAL** | Insula_L, Insula_R |
| **DiFuMo (1024)** | Anterior cingulate cortex anterior LH, Anterior insula (multiple subdivisions), Middle frontal gyrus (multiple subdivisions), Superior parietal lobule (multiple subdivisions), Dorsolateral prefrontal cortex LH, Fusiform gyrus RH. *Full region mapping available in the repository.* |

Table 11: Brain regions and atlas label matching for **Article 3 - (worst Dice score)**. The full list is available in the NeuroConText GitHub repository for *PMID 30885230*.

| Source | Sample Regions |
|---|---|
| **Article regions extracted via Mistral LLM** | **Limbic and Subcortical**: Amygdala, Hippocampus (bilateral), Anterior cingulate cortex (ACC), Middle cingulate cortex.<br>**Frontal Regions**: Dorsolateral prefrontal cortex (DLPFC), General prefrontal cortex, Frontal areas.<br>**Parietal Regions**: Intraparietal sulcus (IPS), Paracentral gyrus (bilateral).<br>**Temporal and Occipital Regions**: Visual number form (posterior inferior temporal gyrus), Left temporal gyrus.<br>**Other**: Insula. |
| **AAL** | Amygdala_L, Amygdala_R, Hippocampus_L, Hippocampus_R, Insula_L, Insula_R, Caudate_L, Caudate_R, Cuneus_L, Cuneus_R, Precuneus_L, Precuneus_R, Putamen_L, Putamen_R, Thalamus_L, Thalamus_R |
| **Harvard-Oxford** | Angular Gyrus, Insular Cortex, Lingual Gyrus, Middle Frontal Gyrus, Occipital Fusiform Gyrus, Paracingulate Gyrus, Postcentral Gyrus, Precentral Gyrus, Superior Frontal Gyrus |
| **DiFuMo (1024)** | Amygdala (anterior/posterior), Anterior insula (multiple subdivisions), Caudate (multiple subdivisions), Hippocampus posterior RH, Cuneus (anterior/posterior/superior), Middle and Superior frontal gyri (multiple subdivisions), Angular gyrus, Fusiform gyrus posterior RH. *Full region mapping available in the repository.* |
| **Juelich** | GM Hippocampus (cornu ammonis, dentate gyrus, subiculum), GM Insula (Id1, Ig1, Ig2), GM Hippocampus entorhinal cortex, GM Hippocampus-amygdaloid transition area |
